# Competition Between Cell-Cell and Cell-Substrate Adhesion Determines Epithelial Monolayer Architecture in Culture

**DOI:** 10.1101/2021.09.13.460154

**Authors:** Christian M. Cammarota, Nicole S. Dawney, Qingyuan Jia, Maren M. Jüng, Joseph A. Glichowski, Philip M. Bellomio, Alexander G. Fletcher, Dan T. Bergstralh

## Abstract

Organ surfaces are lined by epithelial monolayers - sheets of cells that are one-cell thick. This architecture underlies tissue function, and its loss is associated with disease, including cancer. Studies of in-plane epithelial cell behaviors show that a developing epithelium behaves as a fluid in respect to the tissue plane, and can therefore readily adapt to varying mechanical influences during morphogenesis. We asked the question of how monolayer architecture is achieved, and whether it demonstrates the same fluid behavior. To address this problem, we cultured MDCK (Madin-Darby Canine Kidney) cell layers at different densities and timepoints and analyzed their architectures using a novel tool, Automated Layer Analysis (ALAn), which we introduce here. Our experimental and theoretical results lead us to propose that epithelial monolayer architecture is governed by a balance of counteracting forces due to cell-cell and cell-substrate adhesion, and that this balance is influenced by cell density. MDCK cells do not undergo obvious rearrangement along the apical-basal axis; instead, cells that do not contact the substrate aggregate on top of the monolayer. Our findings therefore imply that monolayered architecture is under more rigid control than planar tissue shape in epithelia.

## Introduction

Epithelial tissues line the boundaries of organs, where they perform functions including secretion, absorption, and protection. These functions rely on tissue structure, which is determined in part by the arrangement of the component cells. With respect to the tissue plane, this arrangement has long been considered; D’Arcy Thompson suggested that cells in a tissue would tend to take on shapes that minimize surface contact, in the manner of soap bubbles, in 1917 (1). The soap bubble analogy anticipated the possibility that the arrangement of cells can be dynamic; epithelial cells frequently undergo in-plane rearrangements as a tissue develops (reviewed in (2)). This and other in-plane cell behaviors are well-studied for their contribution to morphogenesis (reviewed in (3)). Consistent with these dynamic cell behaviors, a large body of work shows that epithelial tissues can behave as viscous fluids, and that this fluidity promotes morphogenesis by allowing the tissue to respond to mechanical influence (4–6). As they achieve homeostasis, epithelia can change their material phase, transitioning from fluidity to rigidity (also called unjammed to jammed) (reviewed in (7, 8)).

The descriptions of unjammed and jammed epithelia are based on behaviors that occur in the tissue plane as viewed from above; it is less clear how these different behaviors manifest with respect to tissue depth (the apical-basal axis). In comparison to the tissue plane, the apical-basal arrangement of cells in a simple monolayered epithelium appears straightforward. Cells that comprise a simple monolayer are regular in height, arranged side-by-side, and polarized, meaning that they demonstrate molecular and structural asymmetries. However, while cell arrangement in the apical-basal axis appears less complex, it is at least as important to tissue function. Epithelial disorganization (dysplasia) in this axis underlies disease, including tufting enteropathy, and is a hallmark feature of carcinoma (9). In this study we addressed the question of how monolayered architecture is achieved.

Clues to understanding the development of monolayer architecture are provided by previous work describing the challenges to maintaining it. One route to tissue disorganization is overcrowding, which would be expected to cause “piling up.” Epithelial tissues avoid overcrowding by actively regulating their density. Many cell types exhibit “contact inhibition”, meaning that they downregulate proliferation upon reaching confluence (10, 11). Impairment of this mechanism leads to overproliferation, which has long been associated with dysplasia. In highly crowded areas, homeostatic density can also be restored by active extrusion of live cells (12). Evidence for the importance of this mechanism to monolayer architecture is that impairment (through disruption of the regulator Piezo1) causes the appearance of cell aggregates (12).

Overproliferation is often associated with another route to tissue disorganization, namely the loss of cortical polarity (13, 14). Genetic removal of cortical polarity determinants causes increased cell proliferation in flies and vertebrate cultured cells (reviewed in (15–18)). This is likely because apical and basolateral cortical polarity factors interact with multiple signaling networks that regulate cell proliferation, including the conserved Hippo signaling pathway (16,19–24).

A third potential route towards tissue disorganization is the misplacement of newly-born daughter cells. Daughter cell position is initially determined by the orientation of the metaphase spindle, which sets up the direction of division and is typically aligned in epithelia so that both daughter cells appear within the tissue plane (reviewed in (25, 26)). Misorientation of the spindle can cause a daughter cell to be positioned outside the monolayer, but this does not lead to tissue disorganization. In the *Drosophila* imaginal wing disc, which is pseudostratified, the misplaced cell can undergo delamination and apoptosis (27). This tissue appears to be exceptional, however, since misplaced cells in other tissues simply reintegrate into the monolayer (28–31).

The relevance of material phase (jammed and unjammed) to monolayer architecture has not been addressed. Reintegration might be thought of as an orthogonal example of epithelial cell rearrangement, which is a common in-plane feature of the fluid tissue phase associated with morphogenesis . In this analogy, extralayer cells should be reincorporated in unjammed (densifying) tissues whereas extralayer cells would remain mispositioned in jammed (homeostatic) tissues. Testing this possibility required us to develop a new toolset for quantifying the development of monolayer architecture at mesoscale. Our results suggest that densifying epithelia are not fluid in respect to the apical-basal axis, and that monolayered architecture is dictated by a competition between cell-cell and cell-substrate adhesion.

## Results

### A new automated method to quantitatively classify monolayer architecture of cultured cells

Madin Darby Canine Kidney (MDCK) cells are among the premier epithelial cell models. When cultured on collagen, MDCK cells form an epithelial monolayer, with cells that are cuboidal in shape, joined by E-cadherin at cell-cell junctions, and exhibiting apical-basal polarity (32, 33). We set out to characterize the development of monolayer architecture in this system, but manual observation revealed that it can vary substantially within the same culture well (∼100mm^2^) (Supplemental Figure 1). To solve this technical challenge, we developed a custom-built image analysis toolset called Automated Layer Analysis (ALAn), which implements a set of rules to assess monolayer architecture in a region of interest (Supplemental Text). Across our analyses, these regions are ∼300μm^2^.

ALAn applies a set of rules based on actin intensity and the positions of cell nuclei in three dimensions (Supplemental Figure 2). Nuclear positions are used because they are substantially more reliable and technically straightforward to determine than cell positions. As a corollary, ALAn outputs nuclear number as a proxy for cell number. We tested ALAn using a data set that includes cells plated at different initial densities (1, 2, 4, or 6×10^3^ cells/mm^2^) and allowed to develop over different periods of time (8, 16, 24, and 48hrs). Four patterns, corresponding to different layer architectures, emerged. We used the characteristics of each pattern to define categories, which are classified by ALAn without input from the experimenter.

Three categories demonstrate a single sharp peak in the distribution of nuclei near to the layer bottom, consistent with the expected morphology of an organized monolayer (Figure 1A-C) The fourth category, which we call Disorganized, is based on exclusion. Nuclei in these layers do not organize into a discernable monolayer, but are distributed at all Z-axis positions between the top and bottom of the layer, resulting in a broad nuclear peak (Figure 1D).

**Figure 1:**
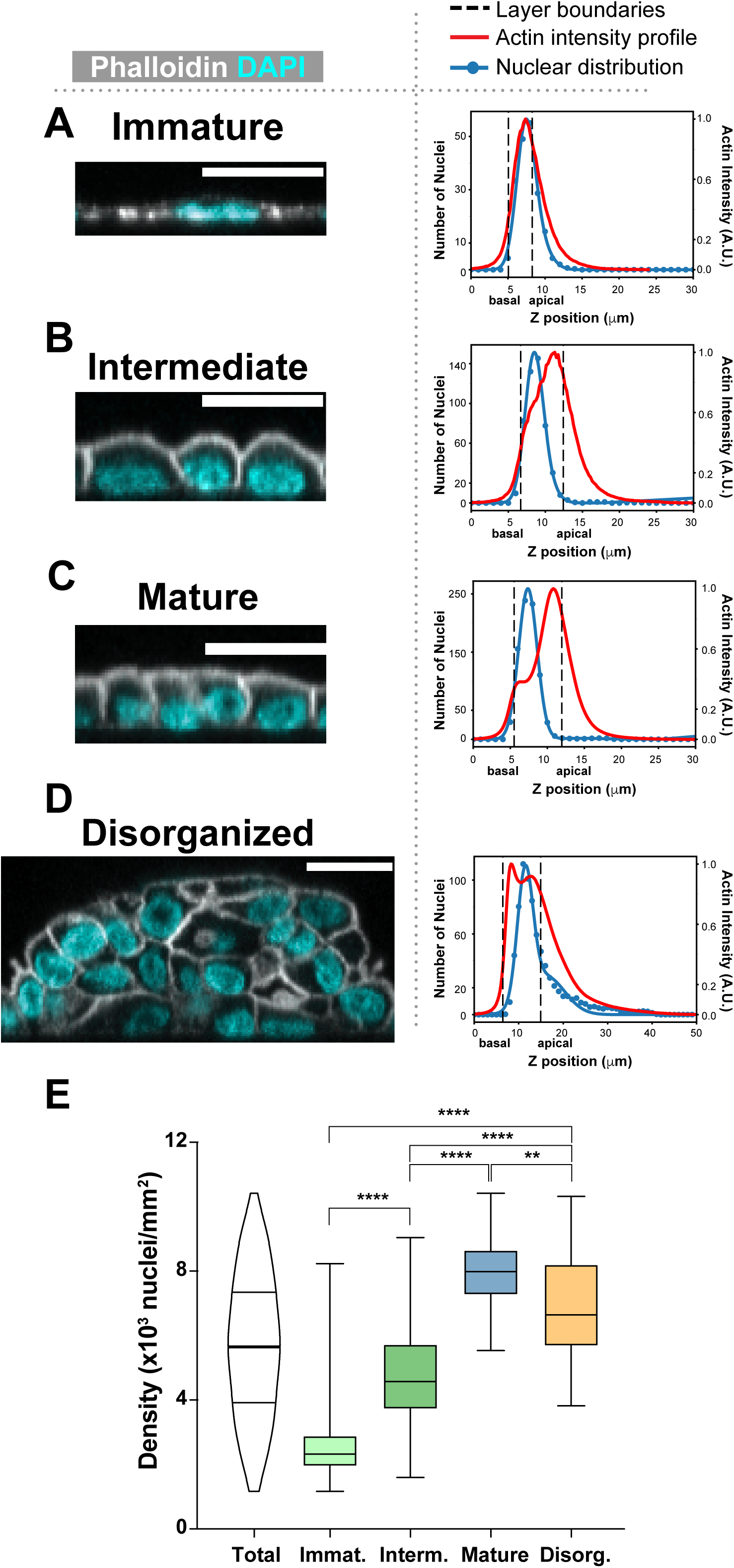
ALAn identifies four distinct layer organizations. **A)** Immature: Component cells of this layer are relatively flat, producing a plot with nuclear centroids above the actin peak. **B)** Intermediate: Component cell have domed apices, shifting the nuclear peak below the actin peak. **C)** Mature: Component cells are regular, with flat apices. A ‘shoulder’ in the actin intensity plot appears. **D)** Disorganized: Nuclei are broadly distributed, as revealed in the histogram of nuclear z-position. **E)** Layer architectures correspond to different cell densities. (Unpaired, two-tailed Student’s t-test).

By their appearance, the first three architectures suggest a developmental series. Cells in the layers we term Immature are short in the apical-basal axis and take on a “fried egg” morphology such that nuclear centroids are positioned at or above the actin intensity peak across the layer (Figure 1A). Cells in the layers we term Intermediate and Mature are generally taller, have a smaller cross-sectional area (in XY), and are more regular (in XY) than cells in the first class (Figure 1B,C; Supplemental Figure 3A-C). Intermediate layer cells are characterized by domed apices, consistent with the cobblestone morphology long described in cultured epithelial models (for example (34–36)). Mature layers are distinguished by the increased length (height) of their cell-cell contacts and a flattening of the apical surface. Across our data set, we found that the three organized layer architectures are separated by density, with some overlap between them (Figure 1E).

Cellular differences are observed in different layer architectures. Consistent with previous work showing that cell-substrate adhesion alone can initiate apical surface identity, we find that cells in all three organized categories demonstrate apical localization of the cortical polarity determinant aPKC (Figure 2A-C, Supplemental Figure 4B) (37). The thick, bright actin signal characteristic of an apical brush-border – which maximizes absorption in the kidney tubule epithelial cells from which MDCK cells are derived - is evident in Mature and Intermediate layers, but less obvious in Immature layers (Figure 2A-C). Furthermore, whereas the adherens junction component E-cadherin is evident in all three layer types, the tight junction marker ZO-1 is clearly defined in Mature layers, less defined in Intermediate layers, and not observed in Immature layers (Figure 2A-C, Supplemental Figure 4D). Together, these results indicate that cell morphology and function are tied to layer architecture.

**Figure 2:**
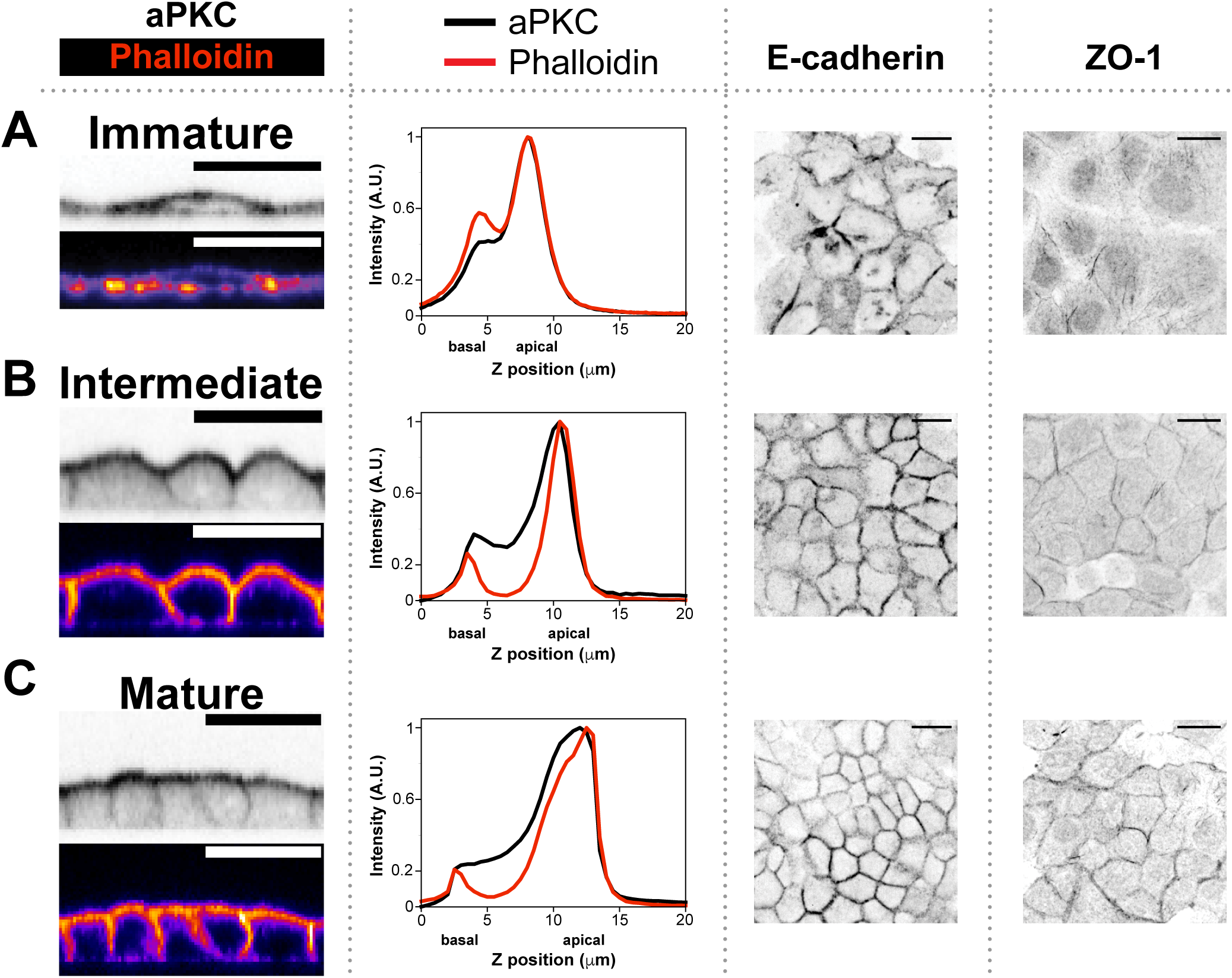
Cell profiles in different organized layer architectures (A: Immature; B: Intermediate, C: Mature). aPKC shows apical localization in all three layer types, and E-Cadherin is observed at all cell-cell borders. Cortical actin shows a two-fold increase at the apical surface in Immature layers and is more asymmetric in the Intermediate and Mature layers. ZO-1 immunoreactivity is not observed in Immature layers, and develops as layers mature. Scale bars = 20µm.

### Architectures transition as layers densify

As predicted by their appearance and associated densities, ALAn revealed that the organized architectures transition over time. Layers with the same initial seeding number (200K cells), shift from predominantly Immature and Intermediate at early timepoints to Intermediate and Mature at later timepoints (Figure 3A). We next sought to distinguish the contributions to architecture made by time spent on the substrate, presumably mediated by integrin signaling, and density, mediated by cell interactions. To do so, we treated cells with the CDK2 inhibitor Roscovitine, which limits proliferation by blocking cell cycle progression at the G1/S transition, thus reducing cell density at a given time after plating (38). Roscovitine-treated cells do not develop the same layer profile over time as DMSO-treated controls; instead they are characterized by both lower densities and shifted towards Immature and Intermediate architectures at the same time point (Figure 3B). We also considered whether the development of the actin brush border was influenced by time spent on the substrate. Cells in Immature layers had the same actin profile regardless of layer age, though a caveat in interpreting this result is that individual cell age is not determined (Supplemental Figure 4C). Together, these results show that time spent on the substrate is insufficient to determine monolayer architecture.

**Figure 3:**
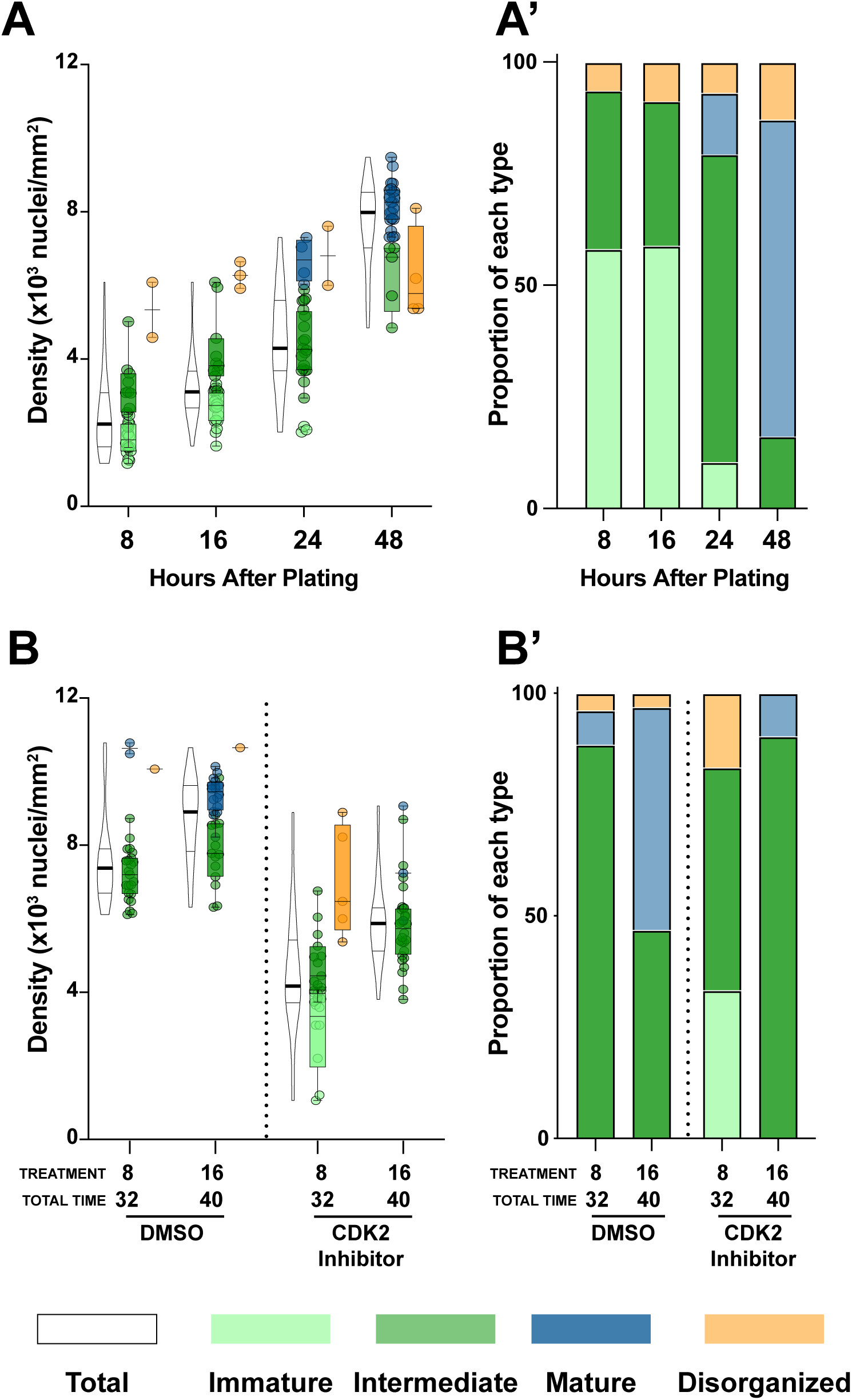
Time spent on the substrate is not sufficient to explain the transition between layer architectures. **A)** Immature and Intermediate layers predominate at early time points, whereas Mature layers are the dominant layer architecture by 48 hours. **B)** Treatment with the CDK inhibitor Roscovitine slows both densification and the maturation of layer architecture. 200K cells were plated in each experiment. Roscovitine was added at 8 or 16 hours and left for 24 hours.

### Intermediate monolayer architecture emerges from both densification and adhesion to the substrate

Our findings thus suggest that layer architecture relies on densification. To explore this possibility, we developed a discrete biophysical model of cell shape dynamics in the apical-basal plane similar to previous ‘deformable polygon’ models (39–41). Cell cortices and the substrate are each discretized into a set of nodes or ‘interaction sites’ which line the cortex of an XZ cellular cross section (Figure 4A). Simulations start with cells positioned above a substrate to recapitulate the starting state of our cultured cells and are run until the cells have reached a steady state. Each cell is subject to the following forces: external adhesions (cell-cell and cell-substrate); internally generated forces regulating size and shape via cytoskeletal dynamics; and an active spreading force driven by actin-based lamellipodia (Figure 4A)(42). External adhesion is modeled by springs that link interaction sites when they come into proximity. The spring breaks when connected nodes move outside of a threshold interaction distance. Internal regulation of cell size and shape is modeled by having each cell maintain a constant area in XZ while minimizing the cortical perimeter. Active cell spreading is simulated in our model with a force (3.2e-5 N, pointing 45° below horizontal) that acts on nodes immediately adjacent to the substrate. Because cells in culture spread actively after an initial passive phase, active spreading in our model is not implemented until 10% of all nodes contact the substrate (42, 43). We modeled contact inhibition of locomotion, which is observed across systems, by turning off the spreading force at each cell side (independently) once that side contacts a neighbor (44). Gravity is estimated to be 4-6 orders of magnitude smaller than the other forces in this system, and is therefore excluded from consideration once cells reach the substrate, as expected in a low Reynolds’ number environment (45). As simulated cells move under the listed forces, there is potential for viscoelastic remodeling; if adjacent nodes are more than twice their resting distance apart along the cell cortex a new node will be created between them, or if adjacent nodes are less than half of their resting length apart one will be removed. The parameter values in our simulation are set by the estimate of ∼100µm^2^ XZ cross section for our MDCK cells, and reported values for physical constants for cultured cells (46–48).

**Figure 4:**
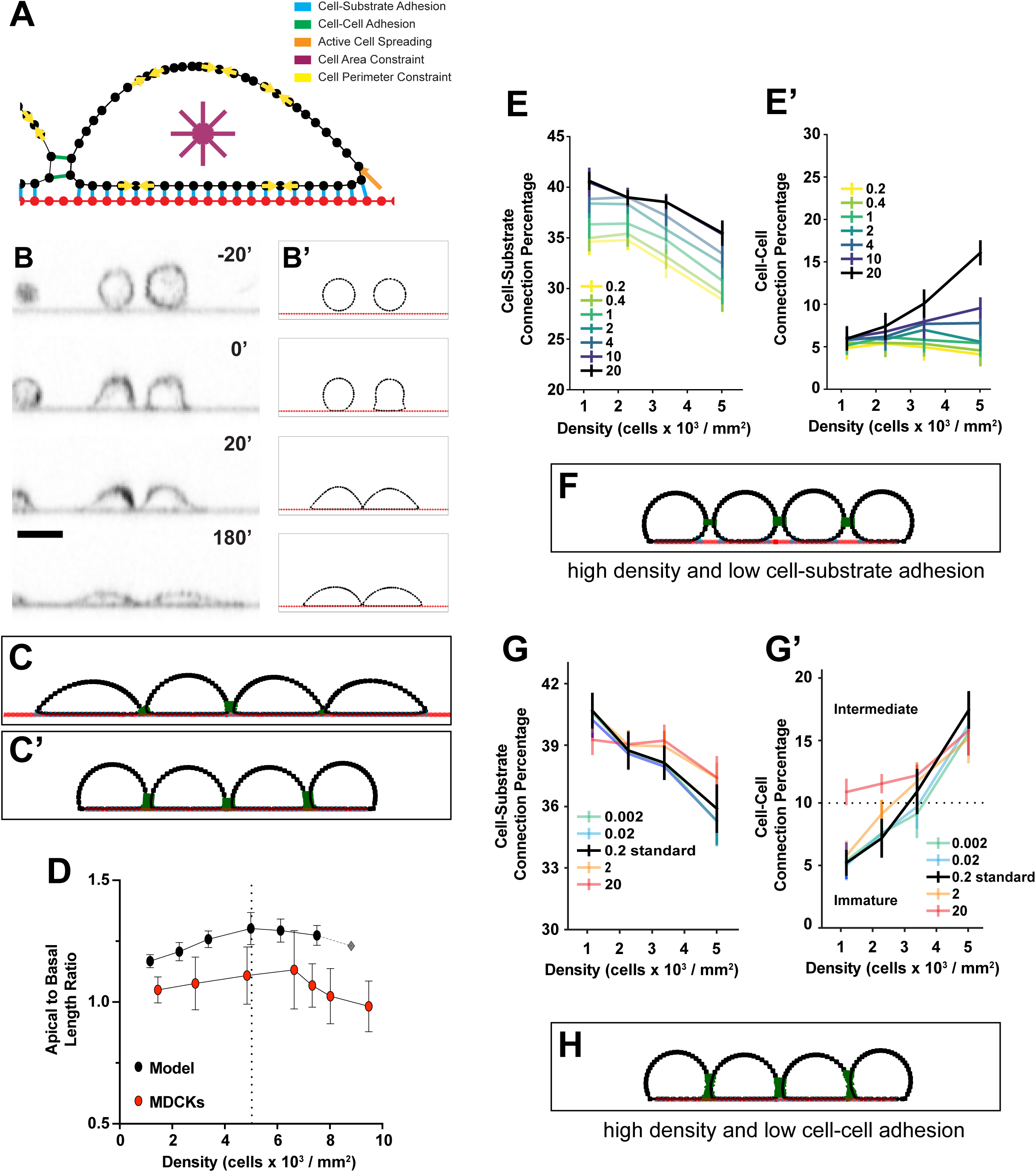
Our viscoelastic cell model recapitulates cell-cell dynamics. **A)** Diagram representing the five forces implemented in the computational model. **B)** Cultured and simulated cells favor spreading over the development of cell-cell contacts (Supplemental Movie 1, Supplemental Movie 2). **C**) Layer architectures in cells modelled at C) low (1.16 x 10^3^ cells/mm^2^) density and normal (*γ*^cell-substrate^ = 20 N/m, *γ*^cell-cell^ = 0.2 N/m) adhesion strengths (Supplemental Movie 3), and C’) higher (4.99 x 10^3^ cells/mm^2^) density and normal (*γ*^cell-substrate^ = 20 N/m, *γ*^cell-cell^ = 0.2 N/m) adhesion strengths (Supplemental Movie 4). **D)** The model predicts a linear increase in apical to basal length ratio at low densities (<5 cells/mm^2^) followed by a linear decrease at higher densities. This ratio is initially similar in cultured and modeled cells, but diverges markedly at mature densities (>6.5 x 10^3^ cells/mm^2^). **E)** As the density of modeled cells increases, the proportion of a cell devoted to substrate connections decreases and the proportion devoted to cell-cell connections increases. The impact of decreasing cell-substrate adhesion strength is shown here. **F)** Layer architecture for cells modelled at high density (7.51 x 10^3^ cells/mm^2^), and low cell-substrate adhesion strength (*γ*^cell-substrate^ = 0.2 N/m, *γ*^cell-cell^ = 0.2 N/m). **G**) The impact of modulating cell-cell adhesion on our simulated cell architectures. Reducing cell-cell adhesion causes Intermediate layers to first appear at higher densities. **H)** Layer architecture for cells modelled at high density (7.51 x 10^3^ cells/mm^2^), and low cell-cell adhesion strength (*γ*^cell-substrate^ = 20 N/m, *γ*^cell-cell^ = 0.002 N/m).

Simulated cells reproduce the dynamic cell morphologies observed in our live imaging (Figure 4B). In both cases, cells tend to maximize substrate connections at the expense of cell-cell adhesions; cells that adhere before contacting the substrate will lose cell-cell adhesions as they spread. Since our model cells do not proliferate, we simulated cell densification by limiting the amount of available substrate. At essentially unlimited substrate, cells spread without developing many contacts (Figure 4C). Lateral surfaces develop as the available substrate is decreased (Figure 4C’, Figure 1B). These surfaces are observed in our model when 10% of all nodes are dedicated to cell-cell adhesion, and we use this point to define the transition to Intermediate architecture. Together, our model and imaging offer a mechanical explanation for the tendency of cells to cover a substrate and reach confluence.

The flat apical surface characteristic of Mature layers is not observed in our simulated cells even at the highest densities, suggesting that the transition from Intermediate to Mature architecture relies on a biophysical mechanism that is not accounted for in our model. To confirm this we compared the ratio of apical length to basal length in MDCK cells and simulated cells. In both MDCKs and simulated cells this ratio increases gradually over the densities associated with Immature and Intermediate architectures. (Figure 1E, 4D). In the simulated cells, a gradual decline begins at a density near 5.5×10^3^ cells/mm^2^ (Figure 1D). In MDCK cells the ratio demonstrates a steeper decline beginning at ∼7×10^3^ cells/mm^2^, correlating with the appearance of flattened apices (Figure 1D). Taken together, these observations indicate that our model recapitulates the development of Immature and Intermediate, but not Mature, architectures. We therefore used it subsequently to model Immature and Intermediate layer development at densities below 5.5×10^3^ cells/mm^2^.

We investigated the importance of spreading behavior to Intermediate architecture in our model by modulating the strength of cell-substrate adhesions. We repeated our simulations over a range of cell-matrix adhesion values from 0.2 N/m, which is the lowest strength that permits spreading, up to 20 N/m, which is physiologically relevant and used as our standard (47). We determined the proportion of total interaction points (per cell) that participate in cell-substrate or cell-cell adhesion, and consider these to be proxies for cell-substrate interface and cell-cell interface length. As expected, we found that reducing cell-substrate adhesion strength reduces the number of cell-substrate adhesions (Figure 4E). Counterintuitively, it also reduces the number of cell-cell adhesions, especially at higher densities (Figure 4F). This is because in our model spreading is outcompeted by the internal regulation of cell shape, which tends towards circularity, at weaker cell-substrate adhesion strengths (Figure 4F).

We also examined the importance of cell-cell adhesion to Intermediate architecture in our model. Starting with our standard cell-cell adhesion strength of 0.2 N/m, which is physiologically relevant, we modulated the strength of cell-cell adhesions in our model by two orders of magnitude in either direction and determined cell-substrate border length and cell-cell border length (48). These parameters normally decrease and increase (respectively) as an MDCK layer matures, and show the same trends in our simulated cells (Figure 4G, Supplemental Figure 5A,B). Modulating cell— cell adhesion strength does not prevent the development of Intermediate architecture in our model (Figure 4G,H). However, it affects the density at which Intermediate architecture is achieved; weakening cell-cell adhesion causes this transition to occur at a higher density (dashed line in Figure 4G), though this effect is modest (3.1×10^3^ cells/mm^2^ in our control versus 3.6×10^3^ cells/mm^2^ when adhesion is reduced to 0.002 N/m). Taken together these results suggest that cell-cell adhesion facilitates the development of Intermediate cell shapes at lower densities (<∼4×10^3^ cells/mm^2^) but is not an absolute requirement.

### E-cadherin is required for Mature, but not Intermediate, monolayer architecture

Our model predicts that the development of Intermediate architecture emerges primarily from densification and spreading. The possibility that cell-cell adhesion has a minor role is unexpected, since MDCK cells A) express the cell-cell adhesion factor E-cadherin, which has long-been associated with epithelial identity and B) are one of the few cultured cell types that effectively epithelialize (Supplemental Figure 5C) (33, 49).

To test the possibility that apical-basal architecture shifts can be independent of cell adhesion, we examined architecture development in HeLa cells, which do not express E-cadherin (Supplemental Figure 5A) (50). We cultured HeLa cells to their maximum density (3.6-4×10^3^ nuclei/mm^2^) using the same collagen-coated culture wells used for our MDCK cell experiments. When adjusted for cell size, this density (4.3-5.2×10^3^ nuclei/mm^2^) falls into a range associated with Immature and Intermediate architectures for MDCK cells. In agreement, ALAn detects these two architectures, as well as some disorganization, in the HeLa cell layers (Figure 5A,B). This result does not reflect a marked change in shape with respect to the tissue surface; even in the Intermediate layers, HeLa cells maintain their characteristic spindle-shaped morphology, which is distinct from the polygonal arrangement of MDCK cells (Figure 5B).

**Figure 5:**
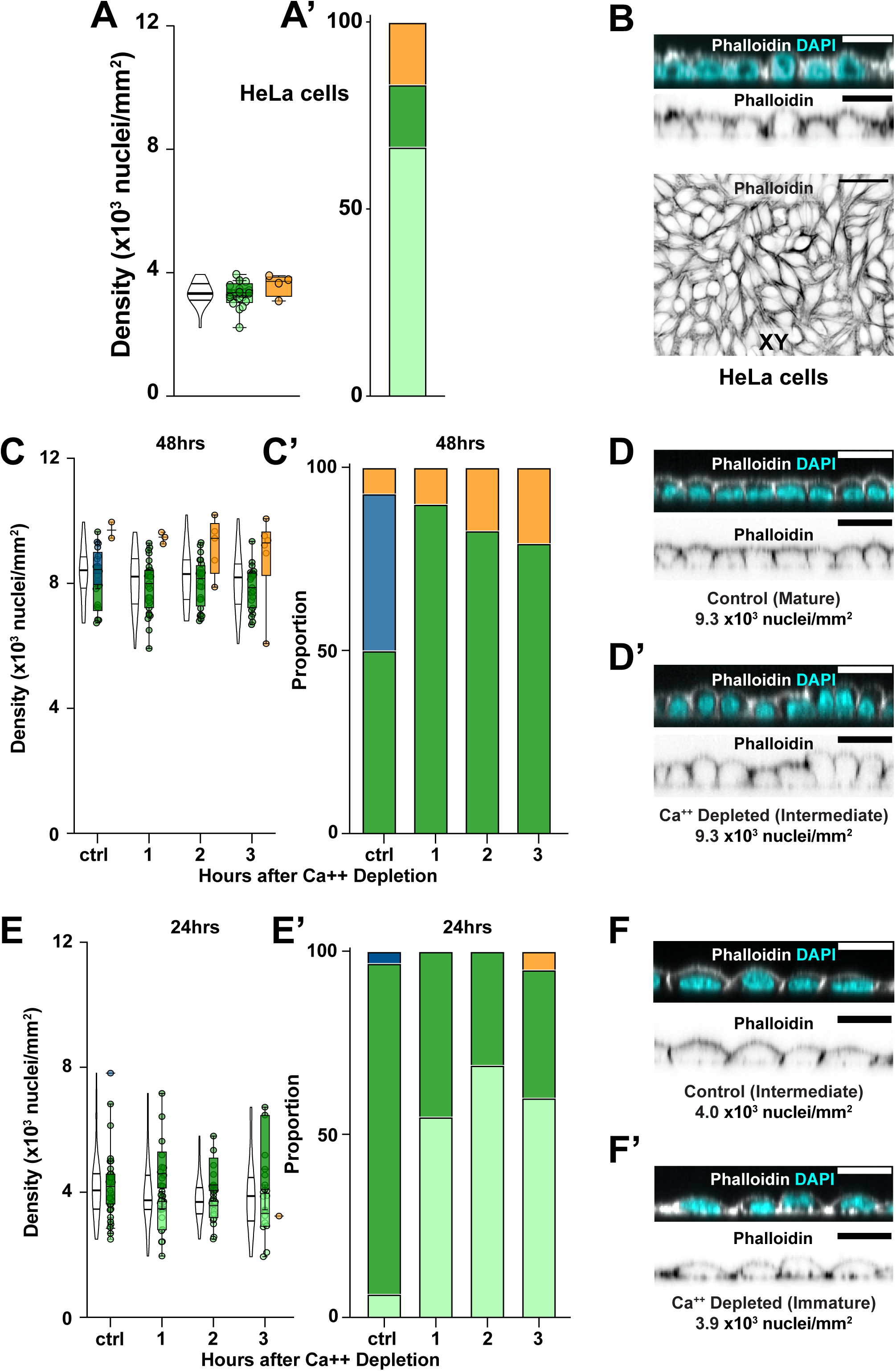
The impact of cell-cell adhesion on layer architecture. **A)** ALAn identifies both Immature and Intermediate layer architectures in HeLa cell culture. **B)** HeLa cells at their maximum density primarily form Immature architectures with a few Intermediate layers (upper). HeLa cells can form cell-cell borders (lower) despite lacking E-cadherin. **C-F)** Calcium depletion causes a change in layer architecture classification. 200K cells were plated. Calcium was removed after 48 hours (C,D) or 24 hours (E,F). **C)** Mature architecture is lost in the absence of calcium. **D)** Calcium depletion causes a change in apical shape at high density. Scale bars = 20µm. **E)** Intermediate architectures are replaced by Immature architectures at densities ≤ 4×10^3^ nuclei/mm^2^. **F)** Calcium depletion causes a reduction in cell-cell border length at low density.

In parallel, we tested the importance of cadherin-mediated cell-cell adhesion in MDCK cells by using calcium depletion to disrupt it at 48 or 24 hours after plating. This manipulation had a strong effect. Calcium depletion results in the disappearance of Mature architectures, as they are shifted to Intermediate even at the highest densities (> 8×10^3^ nuclei/mm^2^) (Figure 5C,D and Figure 1E). This finding suggests that E-cadherin is critical for Mature monolayer architecture. Calcium depletion also results in a shift from Intermediate to Immature architecture, but this shift is only observed at densities ≤ 4×10^3^ nuclei/mm^2^, which is the lowest range for Intermediate architectures (Figure 5E,F and Figure 1E). This observation agrees with our modelling prediction that E-cadherin facilitates Intermediate architecture at low densities, but is not necessary at higher densities. Put another way, we find that the stereotypical cobblestone architecture of epithelial cells in culture can simply emerge as spreading objects crowd together.

### Disorganization arises from a competition between cell-substrate and cell-cell adhesion

Our data and simulations predict that architecture is regulated in two density regimes, and that the second, more dense, of these regimes is associated with a Mature epithelial phenotype. We tested whether high density is sufficient for Mature architecture by examining the development of layer architecture at different initial seeding numbers, using the seeding number of 200K as our standard for comparison. When the seeding number is halved, we find Immature layers predominate at all time points, in agreement with the densities observed (Figure 6A,B). When the seeding number is doubled to 400K, most layers are classified as Intermediate at 8 and 16hrs, while Mature layers develop at later time points and associated higher densities. Again, these architectures agree with the observed cell densities (Figure 6A,B). We further increased the seeding number to 600K, pushing layer densities at the earliest timepoints into the range associated with Mature architectures. However, we find that these layers are not Mature but instead Disorganized (Figure 1E and 6A,B). This finding reveals that density alone does not determine monolayer architecture.

**Figure 6:**
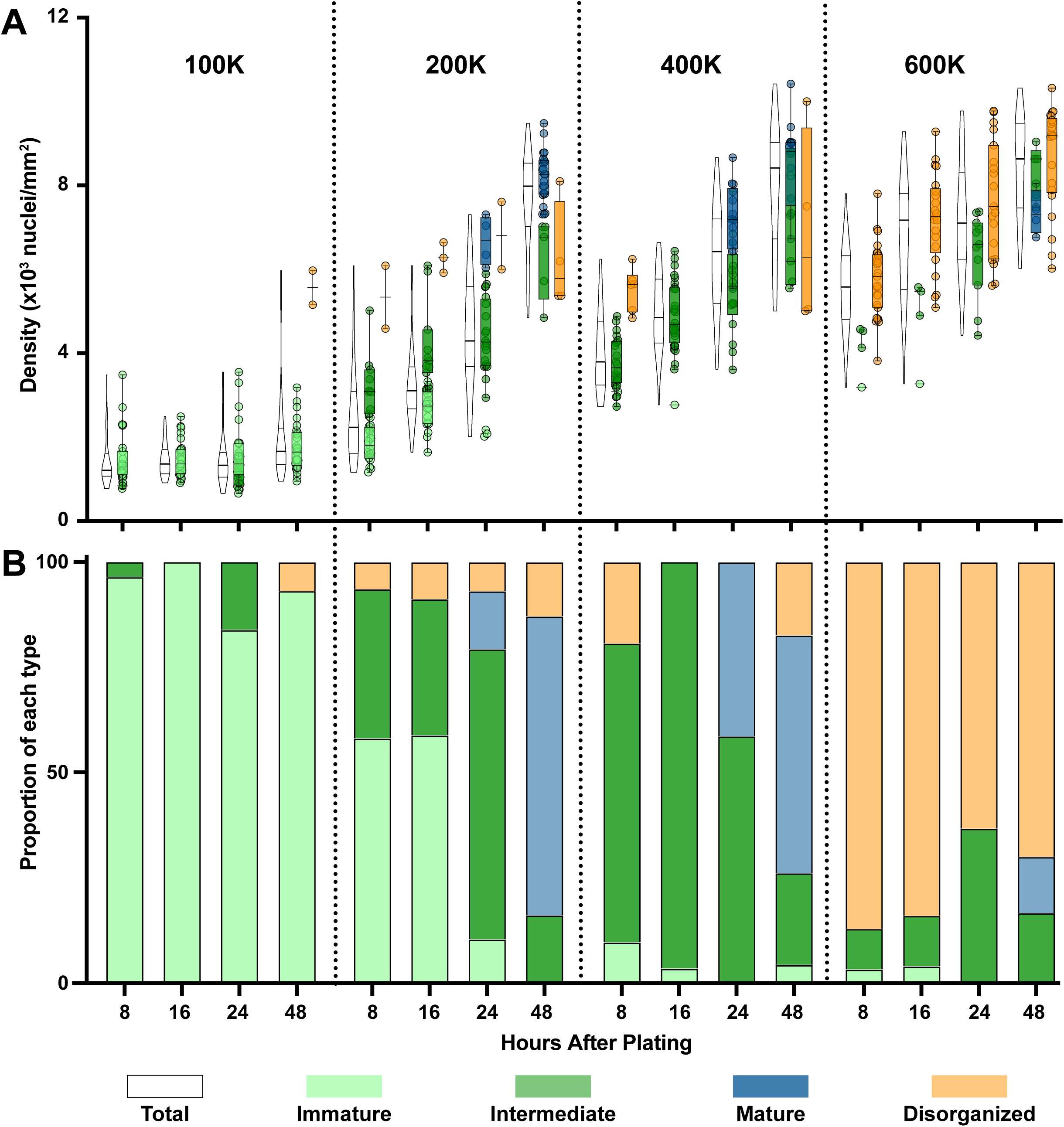
Architecture development at different plating numbers. **A)** Immature and Intermediate layers are observed at a lower densities while Mature layers develop at high densities. Seeding at a high density (600K) results primarily in Disorganization. **B)** Proportions of each layer type found at the four different time points and seeding densities shown in A.

By altering the starting configuration of cells in our simulation, we identified two possibilities that could explain disorganization. The first is that at higher seeding number cells are more likely to encounter each other, and therefore associate into clumps, prior to landing on the substrate (Figure 7A). The second is that the earliest cells to arrive at the substrate, which is limited in area, build a base on which later arriving cells land and attach (Figure 7B). Both possibilities predict cell geometries that appear counterintuitive but are observed in our MDCK cell layers (Figure 7C).

**Figure 7:**
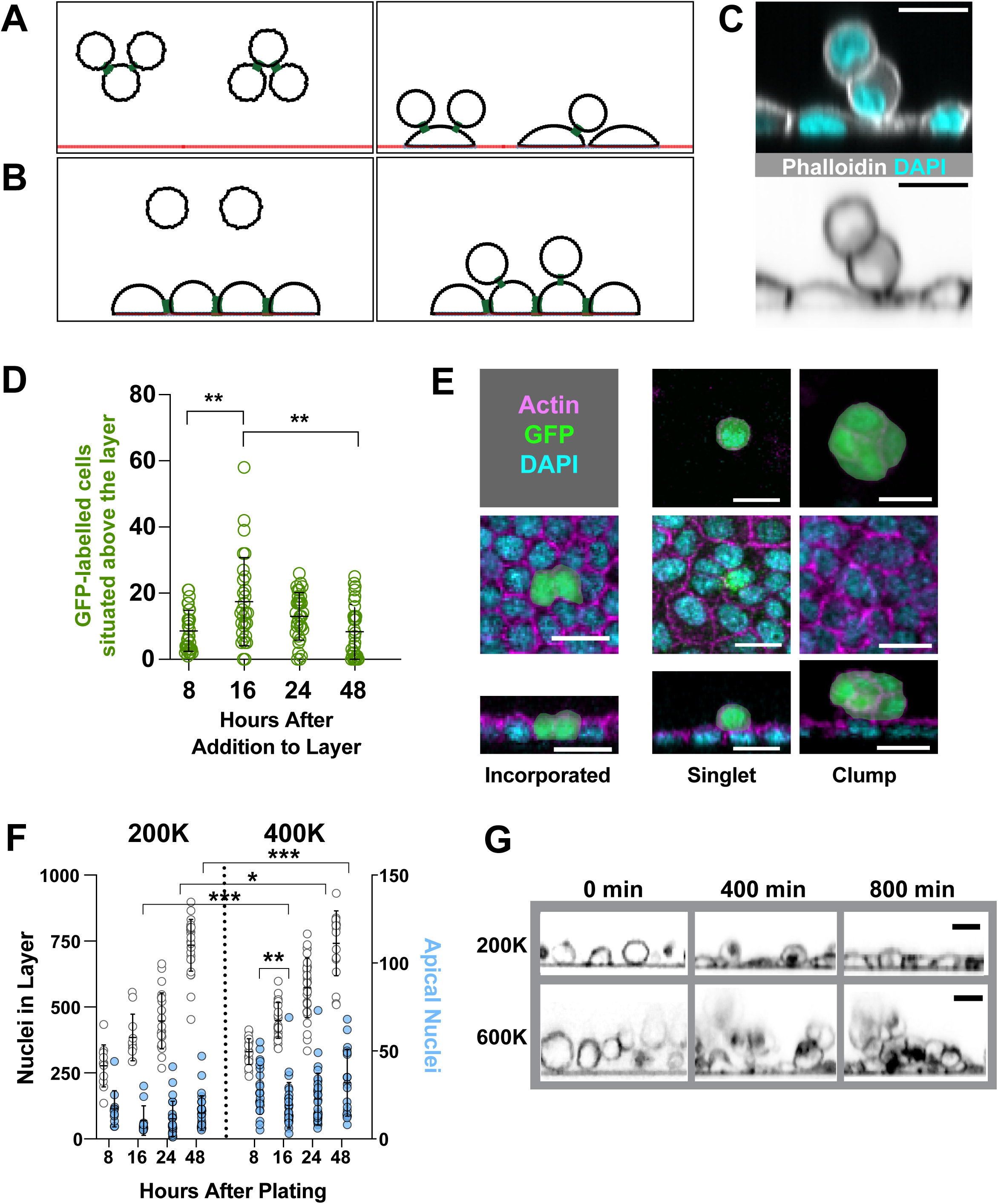
Disorganization arises from a competition between cell-substrate and cell-cell adhesion. **A,B)** Simulating cells dropping in different configurations results in two possibilities to help explain disorganization. Cells can either encounter and adhere to one another before adhering to the substrate (A) or cells that reach the substrate first form attachments and subsequent cells then land on top and attach (B). **C)** Similar configurations are observed in fixed MDCK cell images. **D)** Seeding GFP-labelled MDCK cells on a pre-existing layer shows cells exogenously introduced to a monolayer can attach. **E)** GFP-labelled MDCK cells added to a pre-existing monolayer can be found in different configurations: singlets, clumps or incorporated. **F)** The number of apically-positioned nuclei in Intermediate or Mature layers increases with seeding density. **G)** Live imaging of MDCK cells at low and high plating density (Supplemental Movie 5, Supplemental Movie 6). Cells were imaged for 15 hrs after seeding. Cell membranes are marked with CellMask Orange.

To test whether late-landing cells can stably attach to the layer, we seeded GFP-labelled MDCK cells onto a pre-established monolayer (200K at 24hr, at which point layers are predominantly Intermediate). We found that GFP+ cells attach to the existing layer, peaking in number at ∼16 hours after addition. This number declines gradually over the next 32 hours, suggesting that the GFP+ cells ultimately undergo detachment and/or cell death (Figure 7D). We also considered whether cells undergo rearrangements in the apical-basal axis, and found that this is likely to be rare, since GFP+ cells are seldom incorporated into the layer. However, in contrast to the apically-positioned GFP+ cells, which are frequently observed in disorganized aggregates, the incorporated cells are morphologically indistinct from their non-labelled neighbors (Figure 7E).

Together, these results suggest a “race to the bottom” model in which cell-substrate adhesion, a requirement for layer organization, is in competition with cell-cell adhesion. A prediction of this model is that occasional apically-attached cells should be observed even in organized layers. We tested this possibility using ALAn, which can distinguish and count apically-positioned nuclei. In agreement with our model, we observe apically-positioned nuclei and also find that they increase in number with plating density (Figure 7F). A dip, which is significant at the 400K plating density, is observed between 8 and 16 hours because a small number of cells (∼2%) at the 8 hour timepoint are connected to the substrate but have not yet finished settling down. These cells are indistinguishable from their neighbors by the 16 hour timepoint (Supplemental Figure 6A,B). We also considered whether some of the extralayer nuclei originate inside the layer, perhaps as a result of live cell extrusion in response to increased cell density. While we do not discount this possibility, densification is not the major driver of extra-layer nuclei in our experiments, since their number was not significantly decreased in layers treated with Roscovitine (Supplemental Figure 6C).

Our model is also supported by live imaging of layer development. At lower starting density (200K cells/mm^2^), cells follow a stereotypical pattern: a cell reaches the substrate, undergoes spreading, then develops cell-cell contacts. Live imaging at high starting density (600K cells/mm^2^) reveals cells that remain apical to the underlying layer even up to ∼15 hours, after which point we can no longer image. These extralayer cells adhere to one another to form clusters (Figure 7G). Our results, summarized visually in Figure 8, show that densifying MDCK cell layers do not generally remodel (flatten) in the apical-basal axis, as expected from a fluid.

**Figure 8:**
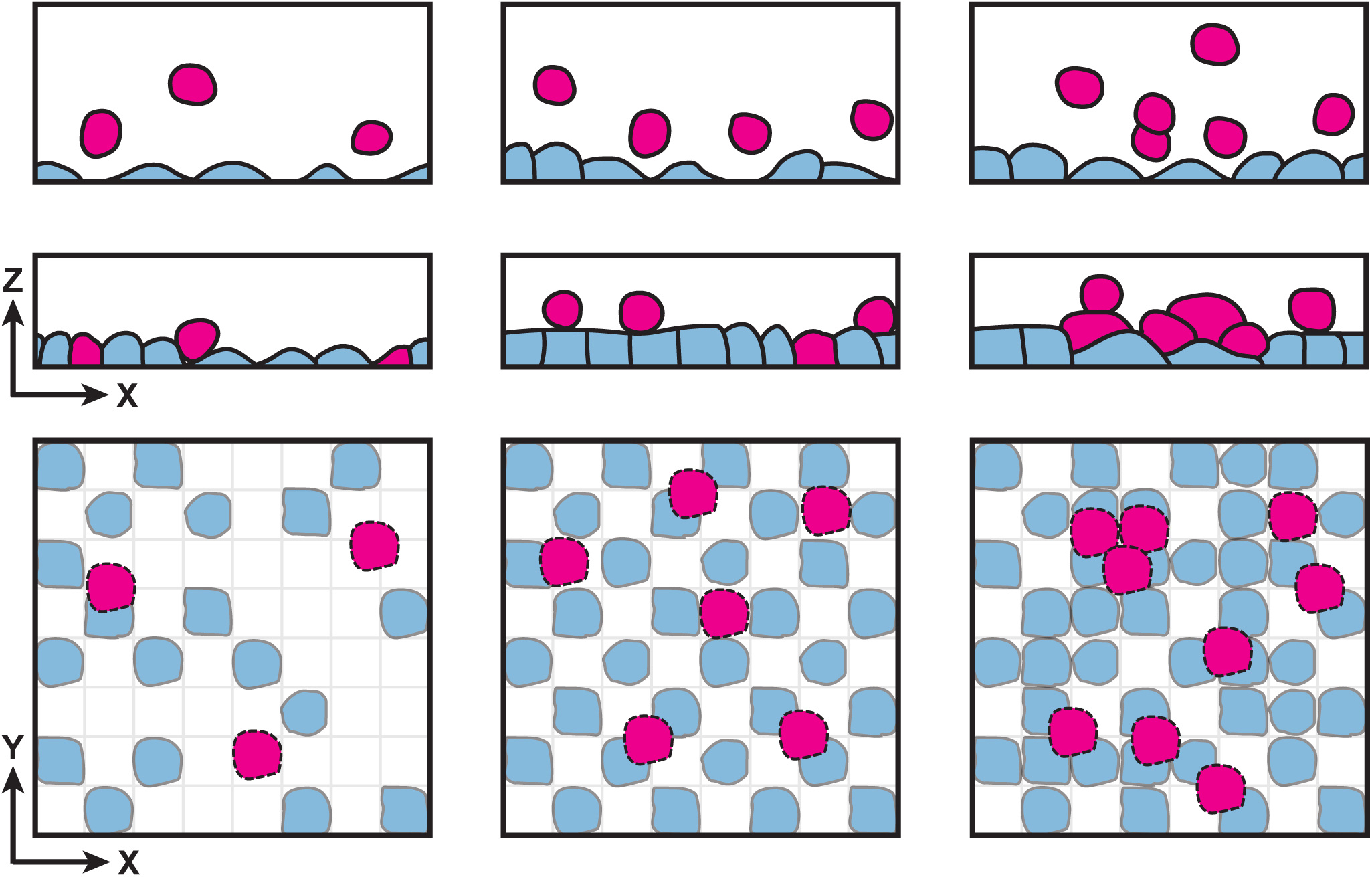
A model for substrate availability vs. seeding density describes the threshold for disorganized plating. Architecture is governed by a balance of cell-cell and cell-substrate adhesion. At a low seeding density, initial cells will adhere to the substrate and spread (blue). Cells that subsequently arrive (pink) have sufficient substrate available and form cell-cell adhesions (left). As seeding density increases, there is less substrate available for the late-landing cells, resulting in an increase in cells apically-positioned on an organized architecture (middle and right).

## Discussion

Our study addressed the question of how monolayer architecture is achieved, and in particular whether the fluid behavior that has been described in respect to the tissue plane extends to the apical-basal axis. We did not find evidence for apical-basal fluidity in proliferating monolayers. Instead, our results suggest that architecture is regulated in two phases, and that these phases are distinguished by cell density. At all densities, architecture is governed by cell-substrate adhesion and the tendency for cells to spread out along a substrate. At lower densities, cell spreading is dominant over cell-cell adhesion, and the first architecture transition (Immature to Intermediate) is dictated by the compression exerted by neighboring cells, with E-cadherin mediated adhesion playing a supplementary role. We speculate that tissue behavior at this point may reflect the physiological importance of an epithelium as a boundary; in other words, the epithelium prioritizes covering a surface over developing a mature monolayer architecture.

The contribution of cell-cell adhesion to architecture increases at higher densities, and is a requirement for the transition to a Mature architecture characterized by flattened apices. One question raised for future study is whether this morphology is important for tissue function. Another is how it is achieved. Though A) external adhesion, B) internal regulation, and C) active spreading are together sufficient to explain the initial appearance of Intermediate architecture, the transition to Mature architecture is not accounted for in our computational model. This may be because our model does not include cadherin-dependent asymmetries in cell mechanical properties that could be important for cell shape. We speculate that the domed appearance of Intermediate apices reflects higher apical surface tension. This tension is increased by apical actomyosin but reduced by the activity of ZO-1 (51–54). Our imaging indicates that ZO-1 is more sharply defined in Mature layers, suggesting the possibility that it is more active in reducing surface tension in those layers. Another potential explanation for the change in apical morphology comes from the recent observation that MDCK cells extend actin protrusions at the apical surface (55). These processes extend into neighboring cells to reinforce damaged adhesions and are suggested to downregulate contractility. The implementation of cortical polarity in our computational model will allow for these asymmetries to be included, and is a goal for future work.

Similar architecture transitions to those observed in our studies take place during the cleavage stages of zebrafish embryogenesis. A series of divisions densifies a developing tissue comprised of blastomeres, a process described as compaction. At the 2-cell stage, adjacent blastomeres do not adhere. At the 8-cell stage, blastomeres take on a cobblestone appearance, with domed apical surfaces, and begin to develop cadherin-based adhesion along their lateral surfaces. Blastomeres at the 32-cell stage are tightly compacted, with mature cadherin adhesions, and demonstrate uniform, flat apical surfaces (56). In light of our cell culture results, these observations suggest that densification promotes epithelialization *in vivo.* This raises the question of whether densification also participates in developmental processes such as mesenchymal to epithelial transition (57).

### Why use ALAn?

Our work shows that obvious experimental parameters, including plating density, confluency, and time spent on the substrate, are imperfect indicators of layer architecture, which can vary substantially across a culture well. Corollary to this, single XZ frames may be insufficient to determine layer architecture at mesoscale. Past studies have had to rely on manual frame selection as a basis for comparison between experimental conditions (for example (58–61)). Here we introduce our image analysis toolset, ALAn, as an improvement over manual determination.

ALAn also facilitates comparison across studies. Our analysis of the literature reveals that MDCK monolayers that we would define as either Intermediate or Mature have been used for studies investigating epithelial architecture (for examples (62, 63)). Given their different morphology and associated cell densities, we speculate that these two architectures have different material properties, and therefore that care must be taken in drawing comparisons between them.

### Epithelia must protect their architecture

Whereas developing epithelia behave as viscous fluids, providing malleability during morphogenesis, we did not find that densifying MDCK layers readily remodel in the apical-basal axis, indicating that this model monolayer behaves as a liquid that cannot be poured. Our findings imply that material properties alone do not protect monolayered architecture from cell misplacement, and therefore that mechanisms for safeguarding it, such as cell reintegration, are active processes.

## Materials and Methods

### Reagents

A list of reagents used in this study can be found in Table 1.

### Cell Culture and Immunostaining

Cells were passaged one day prior to seeding on 8 well collagen-coated slides. Media was changed every 24hrs. After the allotted time period, the media was removed, the cells were washed with dPBS and cells were fixed using ∼4% formaldehyde, 2% PBS-tween for 10 minutes. Three washes (10 minutes each) in PBS-0.2% Tween were carried out between fixation and stainings. Primary antibodies were added at 1:500 dilution as were secondary. FITC phalloidin was used to stain actin. Vectashield plus DAPI was then added to the wells. *Roscovitine –* A concentration of 1.5ug/ml reduced final cell number without obvious toxicity/cell death. *Calcium Depletion -* Media was removed and cells were washed with magnesium-calcium-free PBS. Cells were then incubated for 1, 2 or 3hrs in magnesium-calcium-free PBS with 2mM Magnesium dichloride. *Exogenous cells* - MDCK cells were mosaically transfected using a CMV GFP vector (pLenti CMV GFP Blast (659-1) was a gift from Eric Campeau & Paul Kaufman (Addgene plasmid #17445; http://n2t.net/addgene:17445 ; RRID:Addgene_17445)). GFP positive cells were then selected for and maintained in 10ug/ml blasticidine. 50,000 GFP cells were seeded onto an existing monolayer.

### Imaging and Segmentation

Cells were imaged on an Andor Dragonfly Spinning Disk Confocal microscope using a 40x water objective. Confocal stacks were taken beginning beneath the bottom of the chamber slide where no actin signal could be detected and ending above the layer, when no more actin signal could be detected, taken at a z-spacing of 0.23 µm. To control for differences between culture wells, the same 16 spots were imaged in each well. After image acquisition, the image analysis software Imaris was used to segment all of the nuclei within the images as detailed in ALAn. Seeding number and timepoint were blinded to the experimenter.

Live imaging was performed on a Leica SP5 confocal with a 40x oil objective and images were collected with LAS AF. Z-stacks were taken with at 10 minute intervals, and movies were processed using a Gaussian blur with FIJI. MDCK cells were passaged 24 hrs prior to imaging. The next day, cells were seeded onto collagen coated ibidi slides in a volume of 200ul and immediately stained with 10ul 1:500 dilution of CellMask Orange. A spacing of 1um was used during z-stack acquisition and a stack was taken every 10 minutes overnight.

### Viscoelastic Model

We model each cell as a discrete set of nodes or ‘interaction sites’ which map out the area and perimeter of the XZ plane of the cell and can interact with adjacent cells, or a discretized, linear, rigid substrate. In this model, cells start in a pseudo-random configuration starting with 40 nodes that represent cortical interaction sites which can interact with surrounding objects. We chose to use 40 nodes to give a spacing of roughly 1 node per µm within the XZ plane of a cell but noticed no significant changes to cell dynamics as a function of node spacing. Cells can remodel their cortex, If the spacing between adjacent nodes surpasses 2 times the expected length, a new node will be added at the bisection. If the spacing between adjacent nodes becomes smaller than half the expected length, the node which is closest to its next nearest neighbor will be removed. Remodeling in this way will keep the cells at a roughly constant interaction point density throughout a simulation.

Nodes on a cell undergo both internal regulatory forces and external adhesions with adjacent cells, and the substrate. The internal forces are governed by the following energy equation:

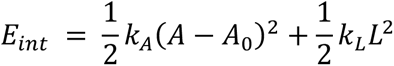

Where *k_A_* controls the constraint on the cross-sectional area of the cell, A is the current cross-sectional area, *A*_0_ is the constant, natural cross-sectional area, *k* controls the constraint on the perimeter of the cell and L is the current perimeter of the cell. Cells tend to round up because of internal regulatory forces in both the model and in culture.

By taking the negative gradient of the above energy with respect to X and Z, we found the internal forces on each node. For ease of calculation, we broke down the energy into its two separate components: 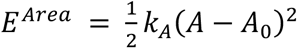 and 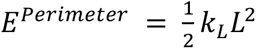. Starting with the shoelace formula for determining the area of a polygon:

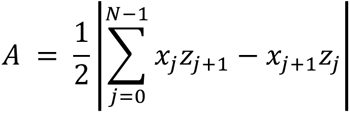

and the perimeter of a polygon:

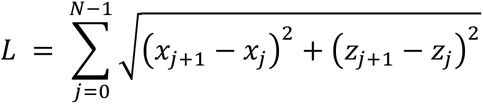

we calculated the forces on each node as a function of the nodal position broken down into the X and Z components. In this description, the N^th^ node is equivalent to the 0^th^ node and the -1^st^ node is equivalent to the (N-1)^th^ node. For the force in the x direction on the i^th^ node of a cell due to the area constraint:

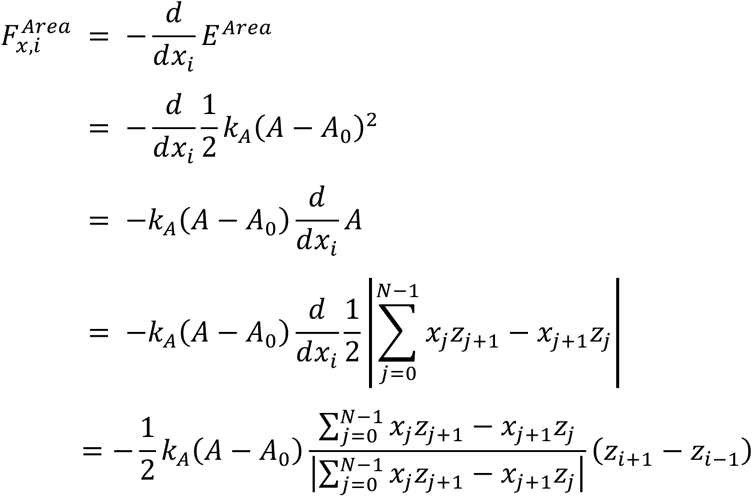

And similarly for the force in the z direction on the i^th^ node of a cell due to the area constraint:

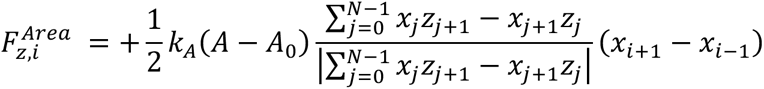

For the force in the x direction on the i^th^ node of a cell due to the perimeter constraint:

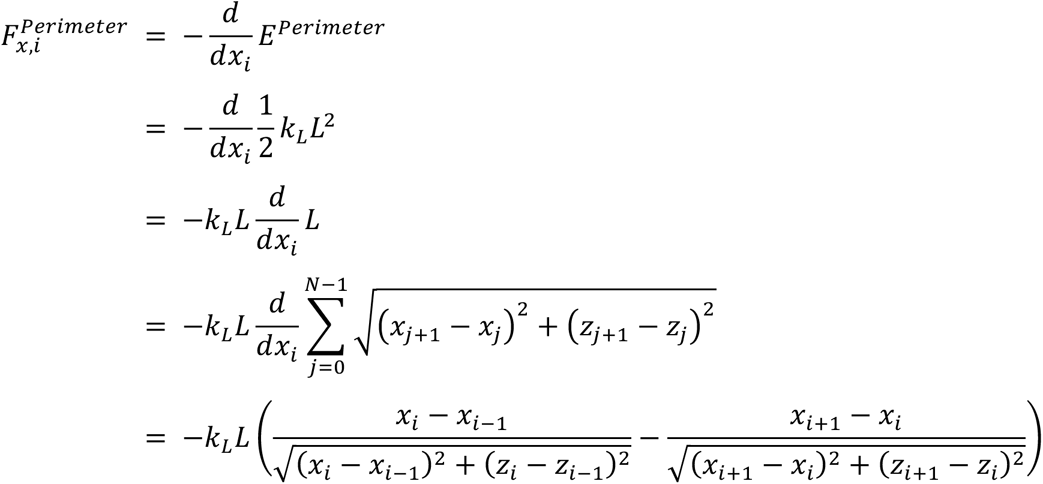

And similarly for the force in the z direction on the i^th^ node of a cell due to the perimeter constraint:

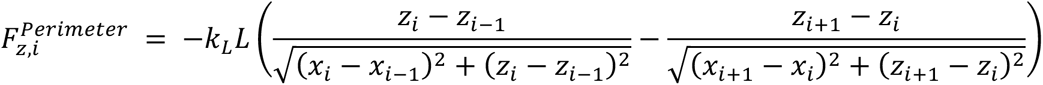

In addition to the internal forces on a cell, cell-substrate and cell-cell adhesions are modeled by Hookean springs with a maximum interaction distance. Cell-substrate adhesions are calculated using:

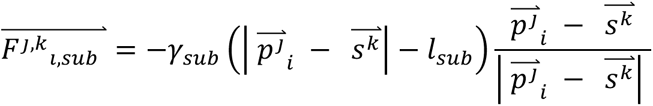

where 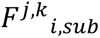 is the force on the j^th^ node of the i^th^ cell due to adhesion to the k^th^ substrate point which points from 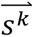 to 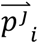 controls the adhesion strength between cells and the substrate, 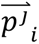 is the position of the j^th^ node of the i^th^ cell, 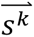 is the position of the k substrate point, and *l_sub_* is the natural length of a cell substrate adhesion. This force only exists when 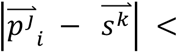 *D_sub_* where *D_sub_* is the maximum interaction distance for a substrate adhesion. Substrate interaction sites will make a connection to only the closest interaction site on a cell, and each cell interaction site can only make one substrate connection.

Cell-cell adhesions are calculated similarly using:

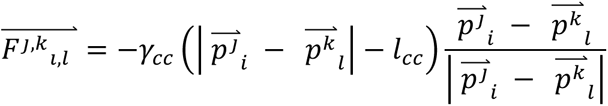

where 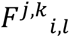 is the force on the j^th^ node of the i^th^ cell due to adhesion to the k^th^ node on the l^th^ cell (which points from 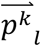 to 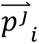), *γ_cc_* controls the adhesion strength between cells, 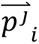 is the position of the j^th^ node of the i^th^ cell, 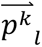 is the position of the k^th^ node of the l^th^ cell, and *l_cc_* is the natural length of a cell-cell adhesion. This force only exists when 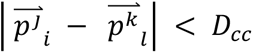 where *D_cc_* is the maximum interaction distance for a cell-cell adhesion.

In addition to the basic physical interactions of the model, two more interactions are implemented to help this model simulate the forces a cell might experience in culture. The first is the addition of gravity so that the cells will tend toward the substrate. Cells exist in a low Reynolds’ number environment, meaning that movement is dominated by viscous forces and that inertial forces are negligible. Gravity is an inertial force and only applies in our system to push the cells toward the substrate at the start of each simulation, and acts on each node. Gravity is set to an arbitrary value of 2e-6 N toward the substrate and is set to 0 N as soon as a cell is within D_sub_ of the substrate, or is connected to such a cell.

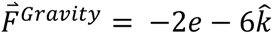

The second implementation is an active cell spreading force. Cell spreading is a two-phase process whereby cells passively spread along a substrate for ∼30 minutes to an hour, followed by an active process likely driven by lamellipodial actin protrusions. To model the active protrusions, a constant force equal to 1.6e-5 N pointing 45° below horizontal acts to the nodes immediately adjacent to the outermost substrate connected nodes.

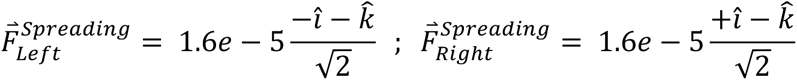

In order to initiate this force, 10% of a cell’s nodes must be connected to the substrate. If there are any cell-cell connections in a space 1/8^th^ of the total number of cell nodes immediately adjacent to a substrate connection, the spreading force will be set to 0 as anticipated by the contact inhibition of locomotion. The left and right spreading edge spreading force is set to 0 independently of one another.

All of the listed forces are summed at each time point to create a net force for each interaction site. This net force is proportional to the instantaneous velocity of each interaction site by a viscous scale factor β:

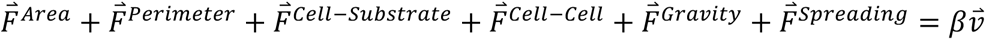

We find the equations of motion for each node using the Euler method of integration from the equation above which relates the instantaneous velocities of each node to their current location. Time is arbitrary in our simulation, and the exact value of the timestep and viscous scale factor are set to β=1 and Δt = 1. We found that at these values, the cells stably change shape/position and reach a steady state shape/position. To time evolve a layer, the first step is the remodeling described above. New nodes are first added, then nodes are removed in all cells synchronously based on the described rules. Then the total force on each node is determined and the new nodal positions are set by the solution to our equation of motion. This model is available on the Bergstralh-Lab GitHub repository: https://github.com/Bergstralh-Lab.

### Estimates of Physical Units from Arbitrary Computational Units

Approximate values for model parameters have been estimated based on current scales for the problem. The model is written with arbitrary units of length, force, and time. Each cell has an arbitrary area of 1 unit^2^. At a cross sectional area of 100 µm^2^ for an XZ cross section of an MDCK cell we get a conversion factor of roughly 10 µm per unit length. This puts our interaction point spacing at roughly 1 µm and the interaction distances at 1-2 µms.

Comparing the simulated cells, which are 2D, with MDCKs, which are 3D, required us to calibrate densities. We first extrapolated our linear density of simulated cells into a rough measurement of the number of cells per square millimeter by assuming the cells to be rotationally symmetric about the z-axis and calculating a planar area. We then set the scale according to measurements for MDCKS: our uncompressed density is set at 1.16 x 10^3^ cells/mm^2^ and the first density at which a 10% cell-cell connection is established is set at 3.38 x 10^3^ cells/mm^2^. Estimations of force from our model are scaled according to published values for cortical stiffness(46). In MDCK cells, the apparent cortical stiffness is 0.1 N/m. The value used for cortical stiffness in the model, k_L_, was 5e-4 force units per length unit. Using the measured value for cortical stiffness and the length unit of the model, the force unit for the model is 2e-3 N. From those scales, we determined the quantities of each of the parameters in our model (Table 2).

### Use of the Viscoelastic Model

To simulate increased cell density, the amount of substrate available to cells was computationally limited. This limitation leads to a scenario where the outermost cells cannot spread further than a certain distance, holding the inner cells at a set maximum density based on available substrate. All measurements at various cell-cell adhesion strength, cell-substrate adhesion strength, and density are a collection of 20 different initial configurations for the cells modelled in each condition. Though all 20 configurations start roughly from the same first line, there are 5 different random seeds used for the pseudo-random initial placing of interaction sites, and 4 different relative intercell spacings for each seed. The measured results are the average of each of the 20 configurations. Each simulation run used 4 cells side by side and the measured quantities were an average of the central two cells. All still images from the model are representative of that density and adhesion strength. There was one instance out of ∼2000 where the simulated cells developed a non-physical overlap due to their initial random placement, and this simulation was excluded.

### Measurement of lateral, apical, and basal surfaces

The segmented line tool in FIJI was used to trace the individual surfaces of cells which were centered in the measured plane of an XZ reconstruction. For the apical to basal length ratio, stills from the cell model were measured similarly in FIJI.

### Actin and aPKC Polarization Ratios

The actin polarization ratio was measured on a per cell basis by first duplicating a circular portion of the z-stack through the center of a cell’s nucleus, making sure not to include cell-cell borders. This cylindrical core was then summed through the XY planes and normalized to create an intensity vs. z plot as done for the entire actin intensity in ALAn. The actin plots made from the cylindrical core have two peaks, corresponding to the apical and basal surfaces (as opposed to ALAn’s plot’s which are primarily one peaked due to the lateral surfaces). The intensity ratio is the normalized apical actin intensity divided by the normalized basal intensity. A ratio of 1 signifies equal apical to basal intensity distribution. A ratio of less than 1 means the basal surface has more actin than the apical surface, and vice versa for a ratio of greater than 1.

## Acknowledgments

We are grateful to Tara Finegan, Holly Lovegrove, the labs of Mark Peifer and Scott Williams, and members of the Bergstralh lab for their questions and comments.

## Author Contributions

DTB and CMC conceived the project. CMC designed the computational model with supervision from AGF. CMC and MMJ coded the model. DTB, CMC, and NSD designed ALAn. CMC, NSD, and JAG coded ALAn. NSD, CMC, QJ, JAG, and PMB performed cultured cell experiments and analysis. DTB, CMC, and NSD wrote the paper.

## Figure Legends

**Supplemental Figure 1:**
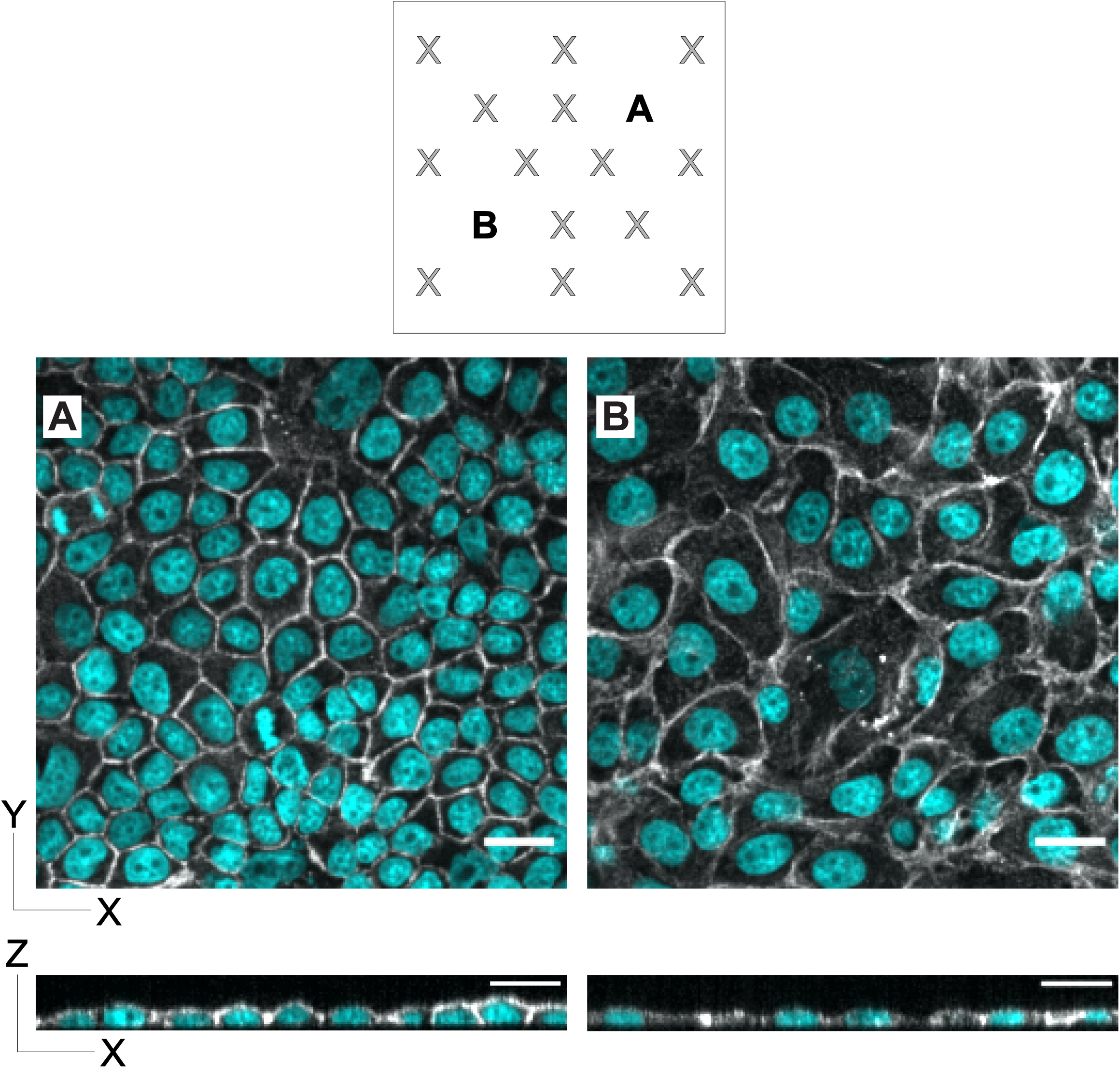
Layer architecture can vary substantially within the same well. **A, B)** The same 16 positions in each well are imaged in our analyses. Here we show layer architectures in two such positions (marked). Cells in A) display a fairly regular planar organization, and have developed lateral borders (bottom). Cells in B) have irregular planar cell shapes and are relatively flat (bottom).

**Supplemental Figure 2:**
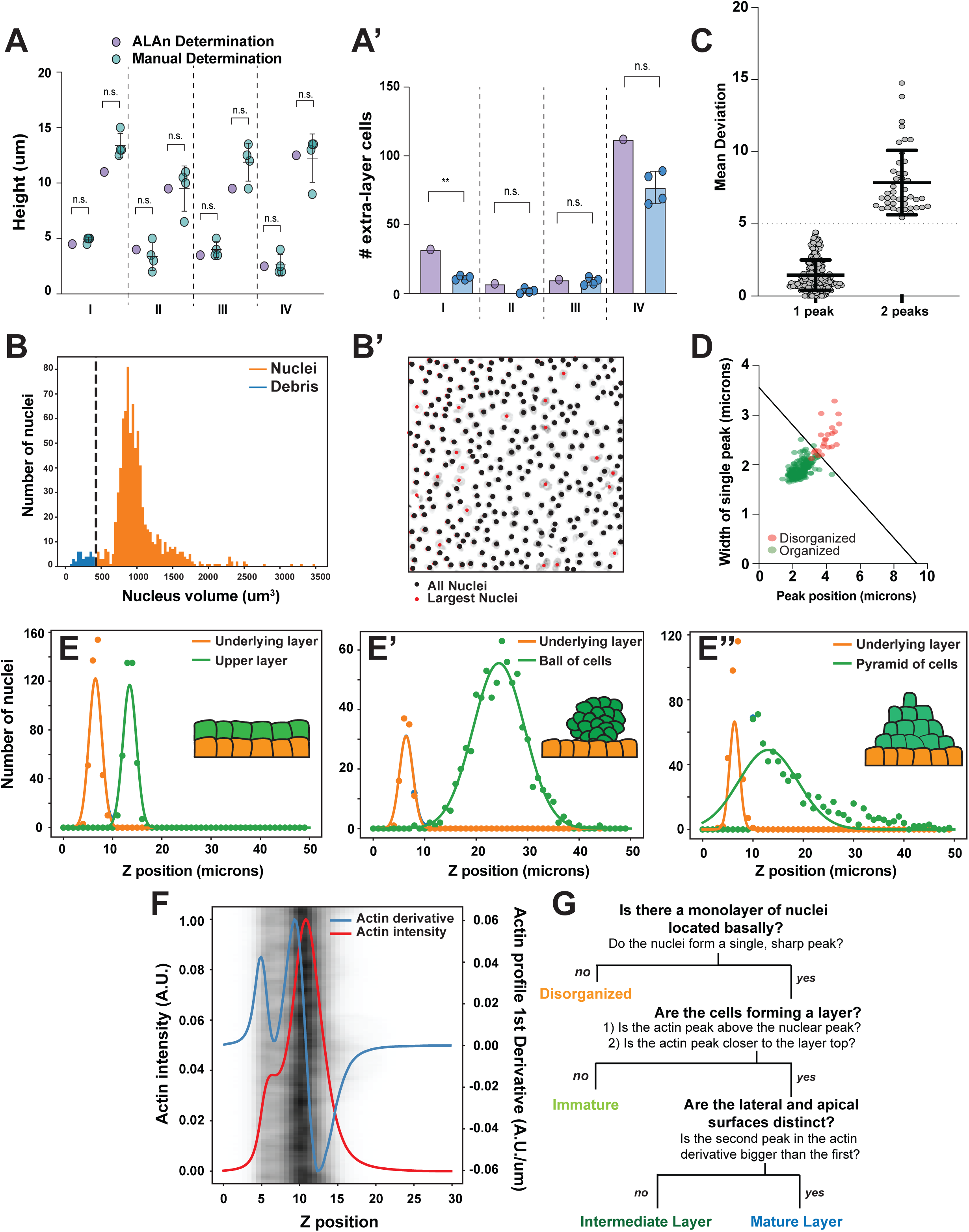
Development and verification of Automated Layer Analysis. **A)** Comparison between manual (performed by 4 researchers independently) and automated determination of A) layer boundaries and A’) extra-layer nuclei. **B)** Segmented objects smaller than 1.5*standard deviation of the mean volume are removed from the analysis. B’) Nuclei are observed above 1.5*standard deviation above the mean volume, so no upper cutoff is set. **C)** A mean deviation cut-off of 5 determines whether the nuclei follow a single (1 peak) or double (2 peak) Gaussian distribution. **D)** If the nuclei follow a single Gaussian distribution, the position of the peak vs. width of the peak determines organization. **E)** Artificial layer examples (E – multilayered; E’ - clump of cells on a layer; E’’ - mountain of cells on a layer) verify that ALAn can accurately determine organization based on the distribution of nuclei. **F)** The first derivative of the actin intensity plot describes the shape of the actin intensity plot and whether one or two peaks are present. **G)** Flow chart depicting the rules set within ALAn to determine each of the four layer types.

**Supplemental Figure 3:**
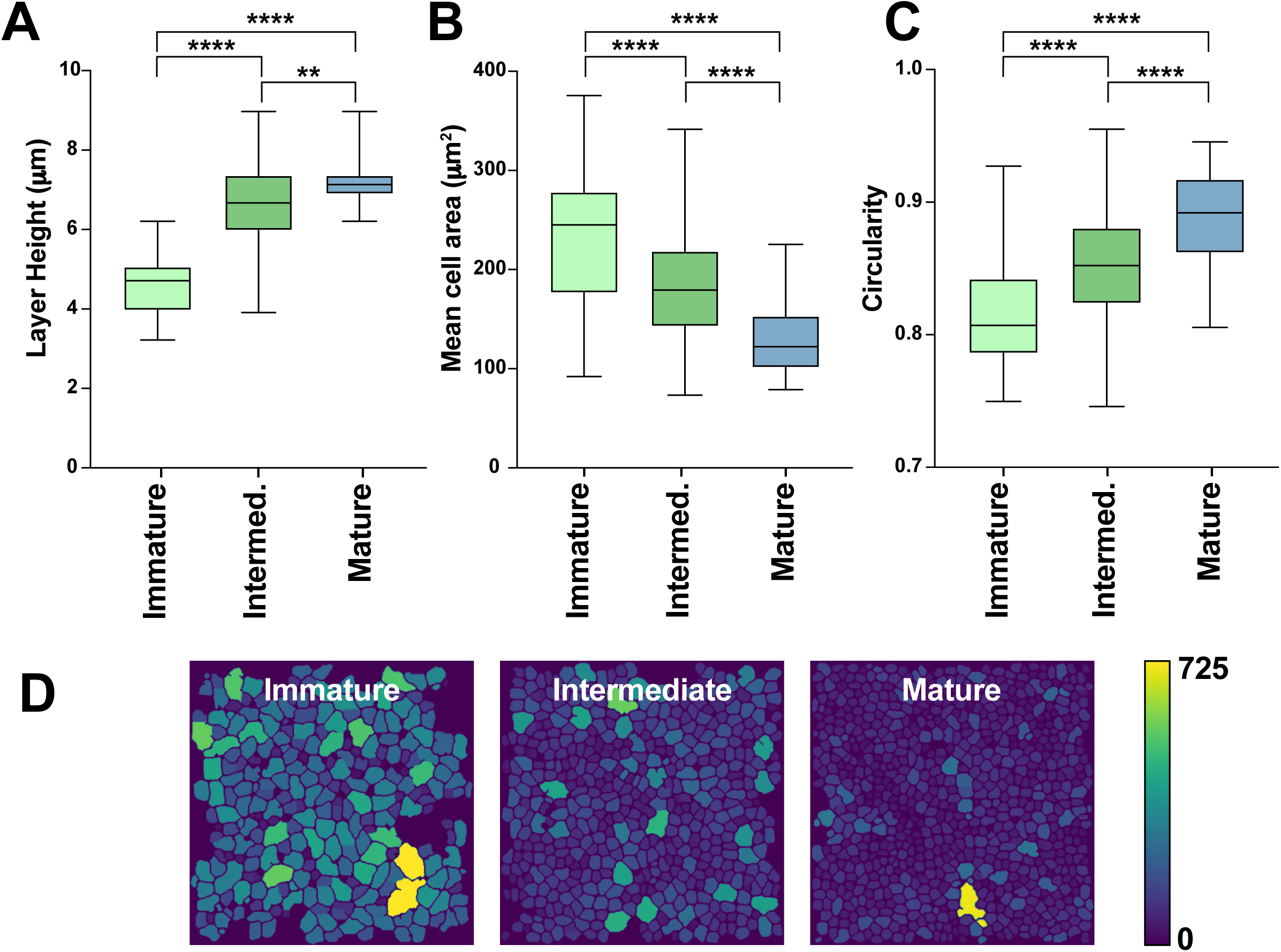
Quantitative description of three layer types. **A)** Height increases as the layer matures. **B)** Cell area decreases as the layer matures. **C)** Circularity increases as the layer matures. **D)** Segmentation of each layer type with a heat map describing cell area. Cells in layers classified as Immature are much larger than those in Intermediate and Mature. Scale in μm^2^.

**Supplemental Figure 4:**
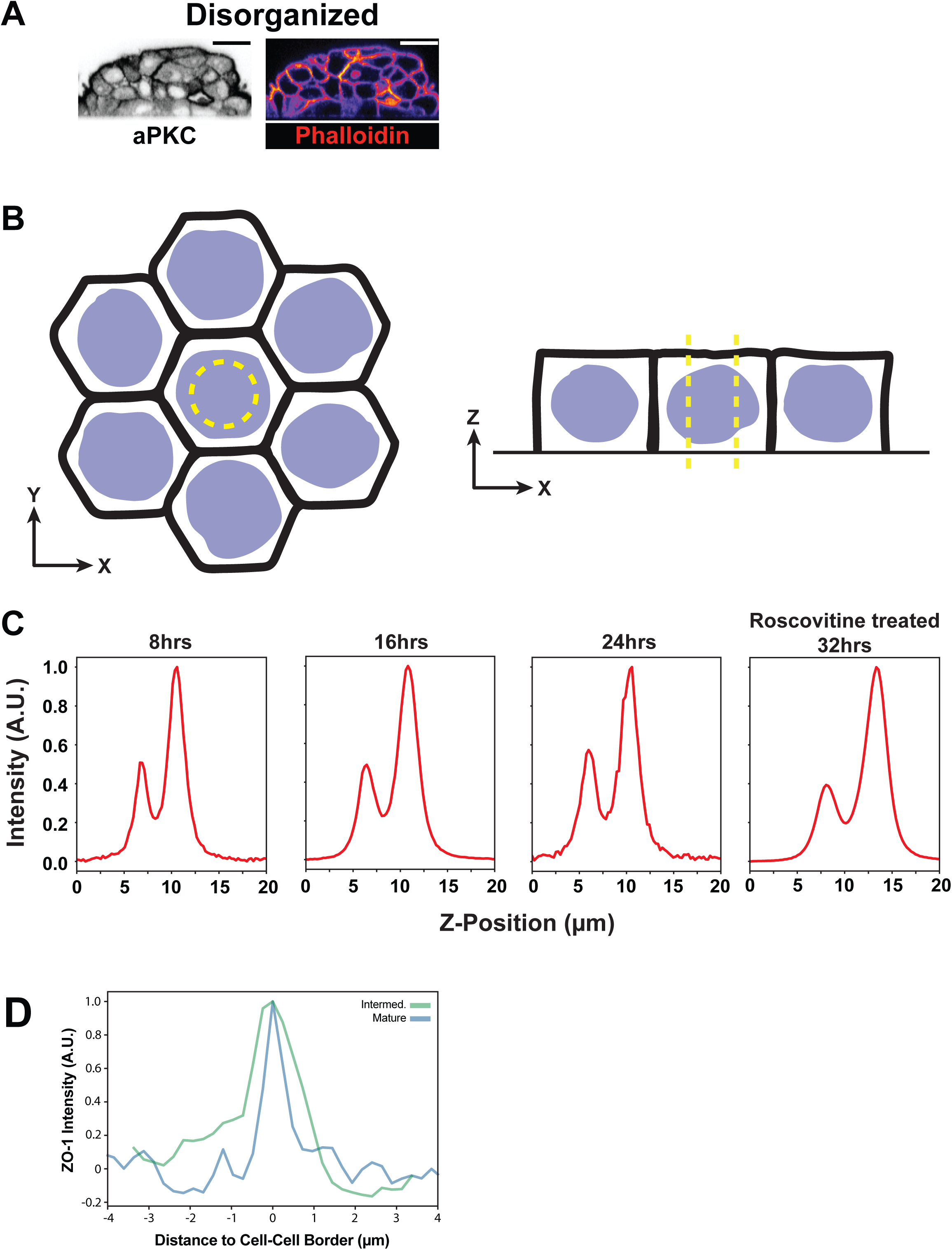
Markers of polarity and adhesion in different layers. A) aPKC is observed at all cell-cell borders in the disorganized regions. **B)** Single cell actin/aPKC plots were made by taking a circular section through center of the nucleus in the XY plane. The X and Y components are summed, resulting in an intensity vs. Z plot which captures apical and basal surfaces while avoiding lateral surfaces. **C)** Cortical actin asymmetry is the same in Immature cells at every timepoint examined. Example plots are shown. **D)** ZO-1 signal at the cell-cell border (position 0) is sharper in a Mature layer cell than an Immature layer cell.

**Supplemental Figure 5:**
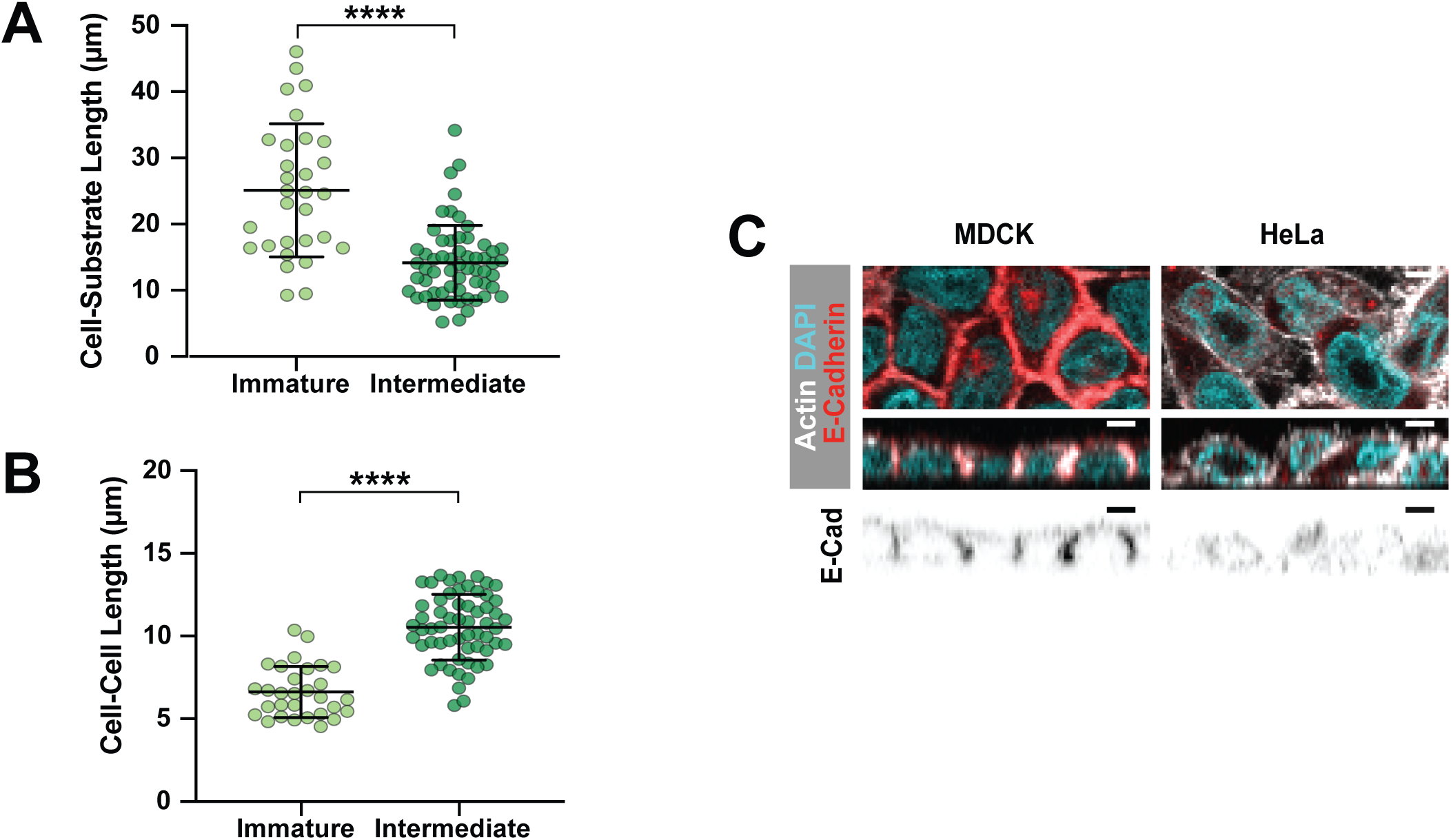
Cell-cell adhesion in MDCKs and HeLas. **A)** The length of MDCK cell-substrate contacts decreases as layers transition from Immature (average cell-substrate length of 25.1 μm) to Intermediate (14.2 μm). **B)** MDCK cell-cell contact length increases as layers transition from Immature (average cell-cell length of 6.6 μm) to Intermediate (10.5 μm). **C)** Unlike MDCK cells, HeLa cells do not express E-cadherin when cultured on collagen coated slides.

**Supplemental Figure 6:**
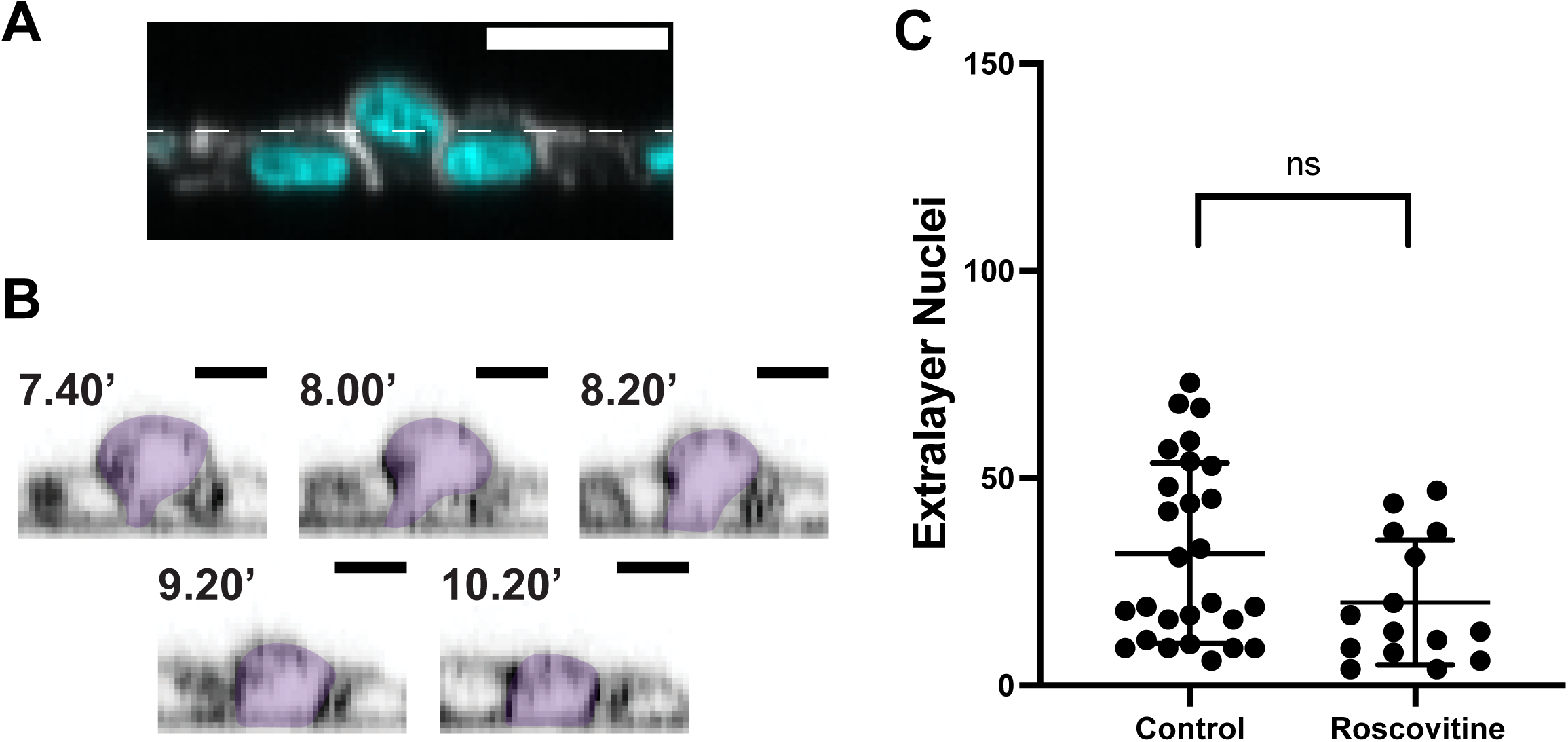
Analysis of extralayer nuclei. A) At 8 hours post seeding, rare (∼2%) cells are attached to the substrate but not yet settled into the layer. The nuclei of these cells can be situated above the rest of the layer. Height of the layer (as determined by ALAn) is shown by the dashed line. B) Example of an apically-protruding cell (false colored) as it finishes settling past 8 hours. C) 200K MDCK cells were plated and allowed to develop for 24hrs, then treated with either DMSO or Roscovitine for a further 24hrs. Comparison between the two conditions shows no significant difference in the number of extralayer cells. Average densities: DMSO control = 9.22 ×10^3^ nuclei/mm^2^; Roscovitine treated = 7.04 ×10^3^ nuclei/mm^2^.

**Supplemental Movie 1:** MDCK cells land on the substrate and spread (Figure 4B). Movie is taken with 10 minutes between frames over the course of ∼4 hours and shown at 10 fps.

**Supplemental Movie 2:** The computational model shows the same cell spreading dynamics as MDCK cells for two isolated cells on a substrate. Movie shows one out of five timesteps up to 12000 iterations. Movie is shown at 100 fps.

**Supplemental Movie 3:** The computational model predicts that four cells prefer to spread along the substrate over making cell-cell contacts. Movie shows one out of five timesteps up to 12000 iterations. Movie is shown at 100 fps.

**Supplemental Movie 4:** The computational model predicts that under artificial densification by substrate limiting, cells will develop and maintain cell-cell contacts. Movie shows one out of five timesteps up to 12000 iterations. Movie is shown at 100 fps.

**Supplemental Movie 5, Supplemental Movie 6:** Seeding different densities of MDCK cells results in different architectures. By 400 minutes, cells plated at 200K condition have spread to form an Intermediate architecture, whereas cells plated at 600K have aggregated to form a Disorganized architecture. Movies are taken with 10 minutes between frames over the course of ∼15 hours and shown at 10 fps.

## Supplemental Text: Development and verification of Automated Layer Analysis (ALAn)

ALAn is a set of image analysis algorithms written in Python, specifically in a Jupyter notebook. ALAn is available in a repository of the Bergstralh-Lab Github: https://github.com/Bergstralh-Lab. It is based on A) the three dimensional distribution of nuclei across a region of interest and B) the one dimensional (Z) distribution of actin in this region.

### Actin Distribution

The actin signal is first projected onto the Z axis by summing the intensity through the X and Y coordinates. The actin intensity plot is normalized by setting the minimum intensity to 0 and scaling the maximum intensity to 1.

### Layer Boundaries

Due to variation in cell size and height across the image, the peak of the intensity plot does not correspond to the top of the layer. We manually determined the focal planes corresponding to the top and bottom of the layer for a subset of images and used these determinations to calibrate the tool. The layer bottom corresponds to the first plane where the actin intensity increases over (0.5-0.2), scaling with density. Likewise, the top of the layer is set to the last plane with an actin intensity above (0.6-0.8). These cutoffs agree with manual determinations performed by four researchers independently (Supplemental Figure 2A). As further evidence that the tool can accurately determine the layer top and bottom, we find that manual counts for number of extra-layer nuclei (also performed by four researchers independently) agree with computational measures (Supplemental Figure 2A’).

### Nuclear Distribution

The image analysis software Imaris is used to segment nuclei based on DAPI signal. In our study, nucleus diameter is set to 7.3µm and smoothed with a filter width of 0.73µm. Nuclei are split by Seed Points and the quality set to 4. Nuclear threshold is set to 120. We remove any objects that are too small to be true nuclei. As segmented nuclear volume can vary with image quality, the cut-off for nuclear volume is set on a per image basis; any volume 1.5 standard deviations below the average volume or lower is excluded (Supplemental Figure 2B). We did not use an upper cutoff, since very large nuclei are occasionally observed in MDCK cell culture (Supplemental Figure 2B’). The distribution of nuclei in the Z dimension is approximated by a histogram with 1 µm bin spacing.

### Layer determination

To determine organization, we fit A) a single Gaussian curve and B) two Gaussian curves to each nuclear distribution, then calculate the average deviation between the fit functions. We do not test directly for multimodality because we do not expect to see two peaks for a disorganized MDCK layer. However, a broad spread of nuclei should be more accurately fit by two Gaussians. If the two Gaussian fit is significantly different from the one Gaussian fit (if the Root Mean Square difference between the fits is greater than 5), the distribution is classed as having two peaks (Supplemental Figure 2C). This method allows us to distinguish between the following architectures:

1. Nuclei in a highly organized layer will either:

a. be fit by a single Gaussian curve peaking at the center of the organized cell layer
b. or be fit by a two-Gaussian distribution with one distinct basal peak, meaning that the overlap of the peaks is <50% of the peak height.
2. If the layer is disorganized, the nuclei will either:

a. be fit by a single, wide (defined as peak width > 3.674 – 0.1819*(the peak location)) (Supplemental Figure 2D). This allows for sparsely packed layers to have more variation in nuclear position, while keeping a strict control on the organization of densely packed layers.
b. or be best fit by a two-Gaussian distribution with overlapping peaks.

We used hypothetical extreme examples of organized layers (Supplemental Figure 2D) to test ALAn as follows.

If a second layer is situated above the first, there will be two sharp Gaussian peaks, corresponding to each layer of cells.
E’) If a large clump of cells sits above the underlying organized layer, the nuclear distribution will demonstrate a sharp basal nuclear peak and a tall, broad Gaussian curve peaking to the right of (above) the first.
E’’) If there is a mountain of cells sitting atop an organized layer, the nuclear distribution will demonstrate a sharp basal nuclear peak and a decaying exponential curve.

Organized layers are then further sub-categorized into Immature, Intermediate or Mature layers. Immature layers are characterized by small or poorly-defined lateral surfaces. Therefore, in these layers the peak nuclear distribution is located at, or occasionally above, the peak actin intensity. Both Intermediate and Mature layers demonstrate defined lateral surfaces. In contrast to Intermediate layer cells, which have a curved apical surface, component cells of Mature layers have flat apices. Therefore, the actin intensity plot in these layers demonstrate a two-step growth to the peak intensity. The first increase (basal actin) is followed by a shoulder (lateral actin), then a second increase from the apical surface (Supplemental Figure 2F). ALAn uses the derivative of the actin intensity plot to detect this shoulder. If the derivative is two-peaked (determined by a prominence of >0.08 using scipy.signal.find_peaks), the ratio of the right peak to the left peak is used to classify maturity. A ratio of greater than one is classed as a Mature layer. Layers for which the ratio is less than one, or for which the derivative has only one peak, are classed as Intermediate.

A flow chart (Supplemental Figure 2G) provides a visual explanation of how ALAn determines layer organization based on the rules outlined above.

**Supplemental Table 1:**
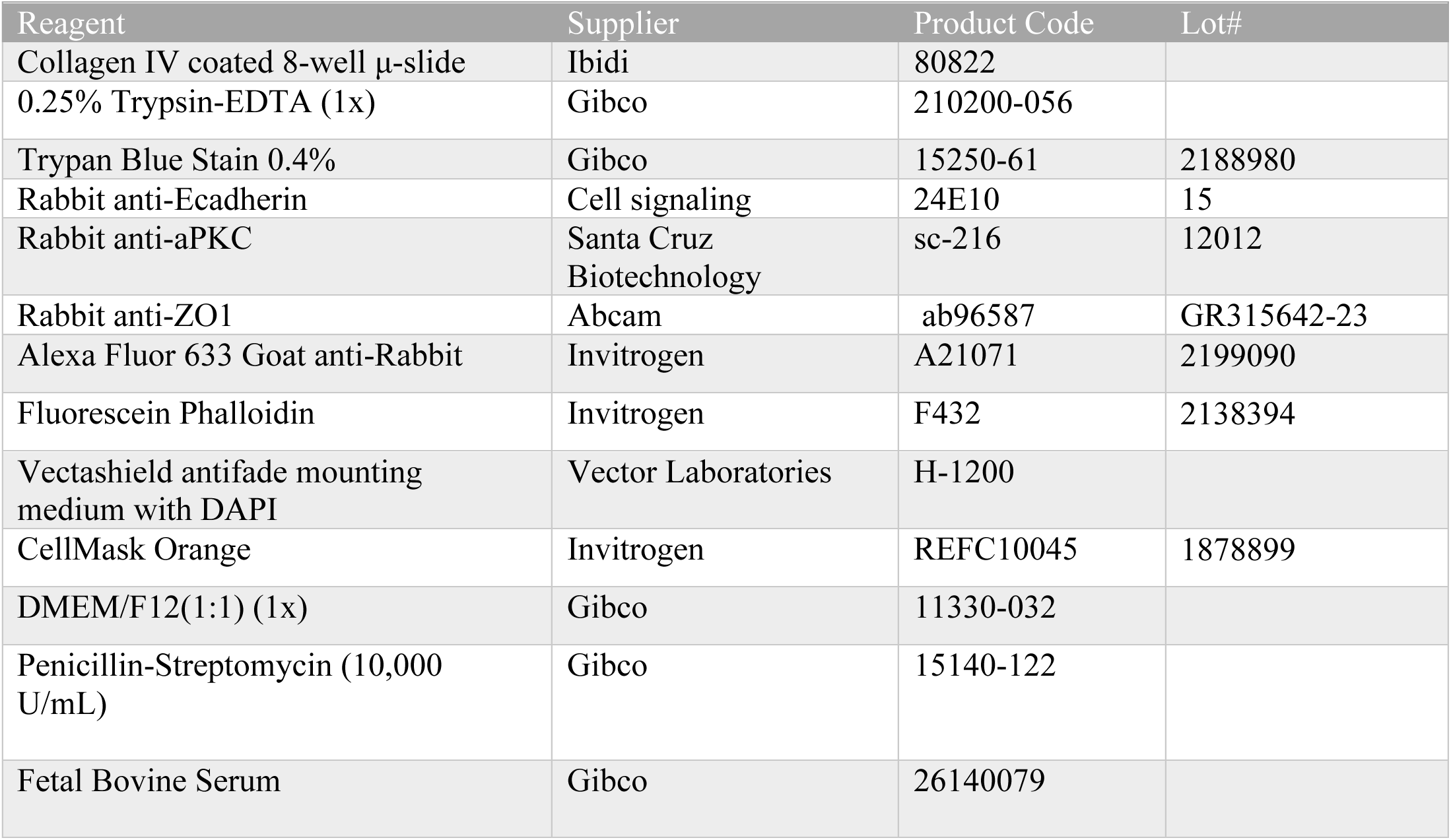
Reagents Used in this Study

**Supplemental Table 1:** Parameters Used in the Viscoelastic Model

| <u>Variables</u> | <u>Arbitrary Value</u> | <u>Physical Value</u> | <u>References</u> |
| --- | --- | --- | --- |
| Cortical Stiffness | 0.0005 (arb) | 0.1 (N/m) | Brückner 2015 |
| Pseudo Bulk Modulus | 1 (arb) | 2e7 (N/m<sup>2</sup>) | n.a. |
| Cell-Cell Adhesion Strength | 0.001 (arb) | 0.2 (N/m) | Nematbakhsh 2017 |
| Cell-Substrate Adhesion Strength | 0.1 (arb) | 20 (N/m) | Gallant 2005 |
| Cell-Cell Interaction Distance | 0.16 (arb) | 1.6 (μm) | n.a. |
| Cell-Substrate Interaction Distance | 0.1 (arb) | 1 (μm) | n.a. |
| Spreading Strength | 0.008 (arb) | 1.6e-5 (N) | n.a. |

